# T helper cell-licensed mast cells promote inflammatory Th17 cells

**DOI:** 10.1101/2021.07.28.454103

**Authors:** Edouard Leveque, Régis Joulia, Camille Petitfils, Xavier Mas-Orea, Gaelle Payros, Camille Laurent, Nicolas Gaudenzio, Gilles Dietrich, Salvatore Valitutti, Nicolas Cenac, Eric Espinosa

**Affiliations:** Inserm, U1037, Cancer Research Center of Toulouse (CRCT), Toulouse, F-31037, France; Toulouse University, Université Paul Sabatier, Toulouse, F-31062, France; National Heart and Lung Institute, Imperial College London, London, SW7 2AZ, UK; Inserm, U1220, Institut de Recherche en Santé Digestive (IRSD), INRA, INP-ENVT, Toulouse, F-31024, France; Department of Pathology, Institut Universitaire du Cancer-Oncopole de Toulouse, CHU Toulouse, 31059 Toulouse, France; Toulouse Institute for Infectious and Inflammatory Diseases (Infinity) INSERM UMR1291 - CNRS UMR5051 - University Toulouse III, Toulouse, France

**Keywords:** Mast cells, helper T cells, prostaglandin E2, IL-1β, Th17, pathogenic Th1.17 cells, Crohn disease, experimental colitis, IL-17, IFN-γ

## Abstract

CD4^+^ T helper cells (Th) infiltrate sites of inflammation and orchestrate the immune response by instructing local leukocytes. Mast cells (MCs) are tissue sentinel cells particularly abundant in skin and mucosa. Here, we analyzed the interplay between human MCs and Th cells and, through the application of RNAseq and functional assays, showed that Th cells induced a specific transcriptomic program in helped MCs (named here MC^TH^) driving them toward an inflammatory phenotype. The gene signature of MC^TH^ indicated that MCs helped by Th cell acquired in turn the capacity to regulate effector T cell response through wide-range of soluble and membrane ligands. Accordingly, we showed that MC^TH^ promoted Th17 cells and notably an inflammatory subset of Th17, producing both IFN-γ and GM-CSF, through a PGE_2_ and IL-1β axis. Our findings demonstrate that activated effector/memory CD4^+^ T cells activate and instruct resting MCs toward a specific differentiated pro-inflammatory phenotype endowed with the capacity to speak back to effector T cells and to mold their functions.

## Introduction

Mast cells (MCs) are tissue resident cells particularly abundant in skin and mucosa. MC response to innate stimuli (complement component C5a, alarmins such as IL-33) or antibody-targeted antigens are well documented (Galli et al., 2020; Halova et al., 2018; Valent et al., 2020). MCs swiftly release numerous prestored mediators by degranulation or *de novo* synthesize cytokines, chemokines and eicosanoids according to their triggered receptors (Espinosa and Valitutti, 2018; Halova et al., 2018) allowing them to participate for instance to inflammatory response initiation, defense against pathogens, venom detoxification, tissue repair or interact with the adaptive immune system most notably CD4^+^ T cells (Dudeck et al., 2019; Valent et al., 2020).

Upon TCR engagement in lymph node, naive CD4^+^ T cells can differentiate into different subsets including the four major T effector (Teff) cell subsets: Th1 (T helper) type 1, Th2, Th17 and T regulatory (Treg) cells according to the signals received during the differentiation process (Ruterbusch et al., 2020; Saravia et al., 2019). CD4^+^ Teff cells then migrate into inflamed tissues where they shape local immune and non-immune cell responses. Understanding how Teff cell functions are induced or shaped in tissues is an area of continuous investigation (Ley, 2014). For instance, local TCR engagement allows Th1 cells to produce cytokines (Honda et al., 2014a; McLachlan et al., 2009), EGFR-expressing Th2 cells infiltrated in the lungs produce IL-13 in response to IL-33 (Minutti et al., 2017) and when exposed to serum amyloid A, Th17 up-regulate IL-17 in the *lamina propria* (Sano et al., 2015). Th cells also show plasticity (*e.g*. enlargement of the panel of cytokines produced) allowing them to adapt and fine-tune the immune response (Mazzoni et al., 2019; Stockinger and Omenetti, 2017; Zielinski et al., 2012).

Subtle combinations of cytokines and environmental cues regulate Th17 plasticity. In response to inflammatory signals, Th17 infiltrating *lamina propria* further differentiate to produce IFN-γ, GM-CSF and IL-17 (Lee et al., 2020; Sano et al., 2015). These potent inflammatory cells required to eliminate pathogens (Acosta-Rodriguez et al., 2007b; Zielinski et al., 2012) may be however detrimental in auto-immune or auto-inflammatory disorders such as multiple sclerosis or inflammatory bowel disease (IBD) (Cerboni et al., 2020; Patel and Kuchroo, 2015; Stockinger and Omenetti, 2017). Nevertheless, environmental mediators promoting pathogenic Th17 cells (referred to as Th1.17) remain poorly understood. We and other have clearly established that MCs can mold CD4^+^ T cell responses by secreting soluble mediators such as TNF or IL-6 (Gaudenzio et al., 2013; Nakae et al., 2005) or by interacting with surface molecules such as OX40L (Kashiwakura et al., 2004). Under certain conditions, human and mouse MCs are able to form immunological synapses with CD4^+^ T cells and promote cytokine secretion (Gaudenzio et al., 2009; Gaudenzio et al., 2013). In turn, Th cells can also trigger the release of mediators by MCs such as oncostatin M upon cell-cell contact (Salamon et al., 2007) or IL-24 by microvesicle secretion (Shefler et al., 2013). The global impact of Th cells on MCs remains however poorly understood.

To better understand Th cell/MC interplay, we examined the profile of primary human MCs following interaction with CD4^+^ T cells using deep RNA-sequencing and mass spectrometry. We found that MCs activated by CD4^+^ T cells (referred to here as MC^TH^) acquired the abilities to release IL-1β and PGE_2_ responsible for the commitment a specific CCR6^+^ CD4^+^ T cell subset toward inflammatory Th1.17 cells. Accordingly, MC-deficient mice were resistant to dextran sulfate sodium (DSS)-induced colitis. In line with a pro-inflammatory role of MCs, analysis of biopsies from Crohn’s patients revealed mucosal MCs expressing both cyclooxygenase 2 (COX-2) and IL-1β. Thus, we identified helped MCs as an original MC functional type able to regulate CD4^+^ T cell-mediated inflammation trough a COX-2/IL-1β axis.

## Results

### CD4^+^ T cell help induces a specific activation program in mast cells

Because tissue resident MCs are expected to encounter infiltrating Ag-experienced CD4^+^ T cells, we incubated human CD4^+^ effector/memory T cells activated or not with anti-CD3/CD28 together with primary human MCs for 24 hours. MCs were then FACS-sorted and processed to mRNA deep sequencing (RNAseq) and genetic profiles were compared with MCs activated by classic MC stimuli (IgE/Ag and IL-33) (Figure 1A). Principal component analysis (PCA) showed that whilst MC co-culture with unstimulated T cells did not impact MC transcriptomic signature, activated effector/memory CD4^+^ T cells induced a clear change in gene expression segregated from the other stimulated conditions (Figure 1B). Proportional-area Venn diagrams depicting the number of differentially expressed genes (DEGs) in stimulated versus unstimulated MCs showed that effector/memory CD4^+^ T cells induced 559 DEGs specific to this activation versus 413 DEGs shared with IL-33 or IgE/Ag or both of them (Figure 1C). The non-supervised clustering of the 500 most significant DEGs clearly gathered IL-33, IgE/Ag and MC+T_ACT_ conditions (Figure S1A). iRegulon, designed to detect transcription factors, targets and motifs/tracks from a set of genes, was used to identify candidate transcription factors that regulate these groups of DEGs (Janky et al., 2014), (http://iregulon.aertslab.org/). Genes upregulated in the 3 conditions were predicted by different sets of transcription factors (Table S1). Interferon-regulatory factor (IRF) protein family were inferred as typical transcription factor of MC^TH^. The analysis highlighted that MCs harbored a specific transcriptomic program following contact with activated CD4^+^ T cells. Of note, gene-expression data obtained by RNA-seq correlated with RT-qPCR measurements (Figure S1B).

**Figure 1.**
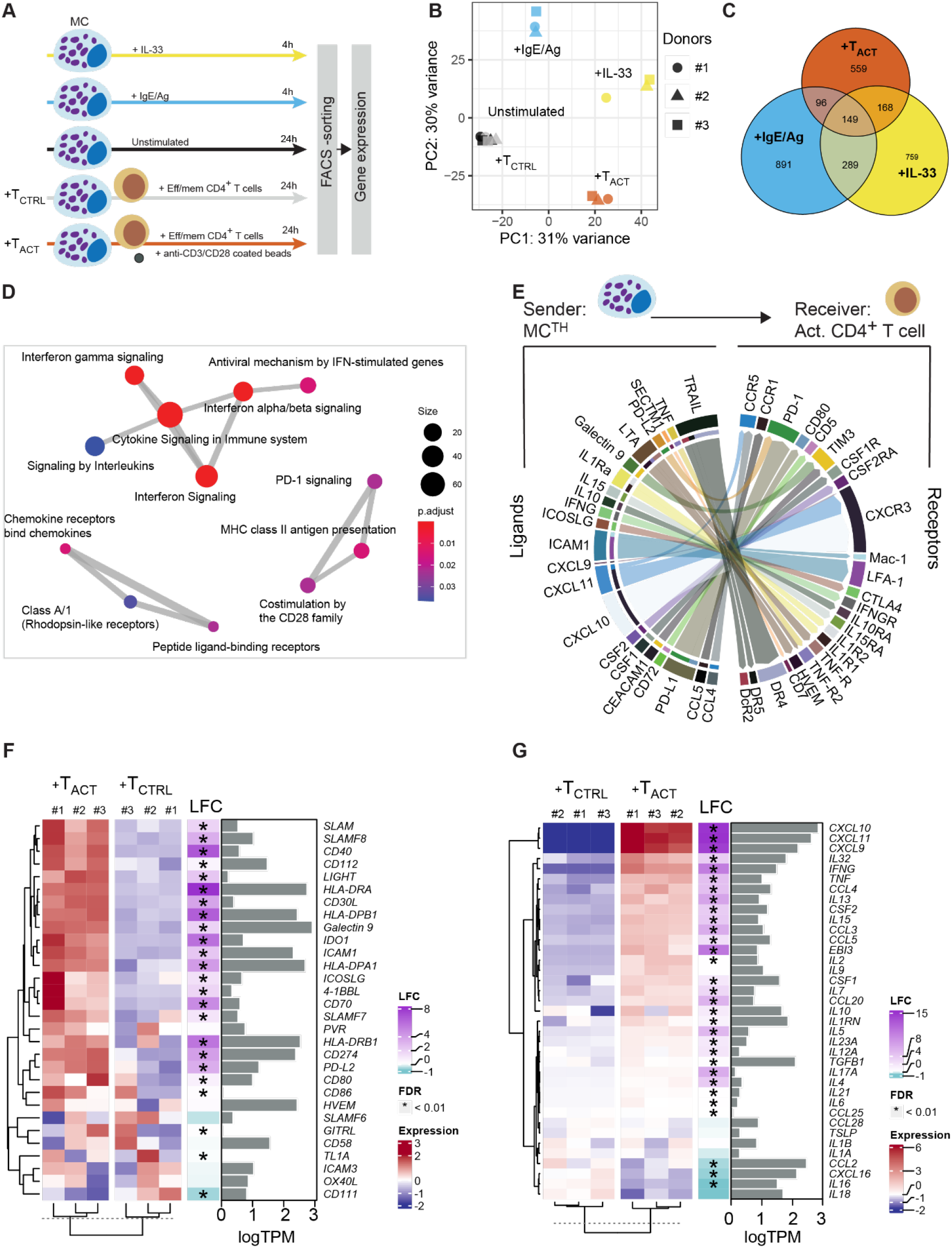
Activated CD4^+^ T cells induce specific gene expression program in MCs. MCs were activated by IL-33 or IgE anti-DNP /DNP-BSA for 4 hours or cocultured with unstimulated (+T_CTRL_) or anti-CD3/CD28 stimulated CD4^+^ effector/memory T cells (+T_ACT_). MCs were next sorted by FACS and subjected to RNA sequencing. (A) Experimental setup. (B) PCA analysis of sample relationship (n=3 donors). The percent of variance explained by each component is reported in parentheses. (C) Proportional-area Venn diagrams showing the overlap of DEGs (absolute log_2_ fold change ≥ 1.58; adjusted p value (FDR) ≤ 0.01) in activated MCs (+IgE/Ag, blue; +IL-33, yellow; +T_ACT_, red) versus resting MCs. (D) Gene set enrichment analysis (Reactome pathways) of MC DEGs (Log_2_FC>1.58 and FDR<0.01) between MC +T_ACT_ vs MC +T_CTRL_ conditions (FDR<0.05). Each node represents one gene set (i.e. a Reactome pathway) where the node size indicates the number of genes enriched in the gene set and the color indicates the padj-value and the edge thickness is proportional to the number of overlapping genes between pathways. (E) Ligand-Receptor interaction inference and visualization. Top 30 strongest predicted interactions visualized by circle plot depicting links between predicted ligands from MC^TH^ (only ligands corresponding to DEGs from MC +T_ACT_ relative to MC +T_CTRL_ were analyzed) with their associated receptors expressed on activated memory CD4^+^ T cells. The arrow thickness is proportional to the computed L-R score. Multiples HLA molecules and their receptors were not displayed. (F-G) Clustered heatmap of genes coding for HLA class II (only genes coding one HLA-DP and one HLA-DR are shown) and co-stimulation molecules (F) or main chemokines and cytokines for which CD4^+^ T cells express the receptor (G). Heatmaps show the relative expression level of each gene between MC +T_ACT_ vs MC +T_CTRL_ conditions. Each gene expression level in +T_ACT_ condition (as depicted by transcript per kilobase million, TPM) and its log_2_-fold-change expression and associated FDR (stars indicate FDR<0.01) are also shown. IL12B was not expressed by MC^TH^ and is not shown in the heatmap. See also figure S1, table S1, S2 and S3.

A differential gene expression analysis of MC^TH^ versus MCs co-cultured with unstimulated CD4^+^ T cells revealed 856 DEGs (Figure S1C and Table S2). Gene ontology analysis using the Reactome database showed that these DEGs mostly pertained to intercellular communication pathways and pointed out some MC^TH^ DEGs included in pathways related to immunological synapse formation (Figure 1D).

Because these results suggested that MC^TH^ increased their ability to communicate with Th cells, we analyzed, based on our transcriptomic dataset, the up-regulation of ligands in MC^TH^ and that of their cognate receptors based on CD3/CD28-activated memory CD4^+^ T cell dataset (LaMere et al., 2017). We computed a communication score for each L-R pair in a publicly available database by adapting the computational method described in the ICELLNET framework (Noel et al., 2021). This score is based on both expression level thresholding method (to determine active L-R pair) and on R-L expression product method (to rank the active L-R pair) (Armingol et al., 2021) with MCs as sender cells and activated Th cells as receiver cells. We found 66 active L-R pairs (Table S3) and mapped the interactions of the top 30 expressed L-R pairs (Figure 1E). L-R pairs referred mostly to cytokine/chemokines (e.g. CXCL9, CXCL10 and TNF) or co-stimulation molecules (e.g. ICOSL, ICAM-1, PD-L1).

To thoroughly analyze the potential of MC^TH^ to provide ligands involved in immunological synapse formation with Th cells, we defined two sets of genes: i) those related to signal 1 and 2 (Chen and Flies, 2013; Sharpe, 2017) ii) those coding for cytokines and chemoattractant molecules for which activated memory CD4^+^ T cells expressed the cognate receptors (from bulk RNA-seq data (Gutierrez-Arcelus et al., 2020; LaMere et al., 2017)). The analysis of the gene expression showed that several genes implicated in the immunological synapse such as HLA-DR and co-stimulation molecules such as CD80, PD-L1 and ICAM-1 were upregulated and expressed with a transcripts per kilobase million (TPM) >10 (Figure 1E). These molecules were increased at the MC^TH^ surface after 48h of coculture (Figure S1D). Likewise, MC^TH^ increased the mRNA levels of chemokines able to attract activated T cells such as CXCL9 and CXCL10 (Figure 1F). Analysis at the protein level showed that MC^TH^ acquired the ability to produce these chemokines (Figure S1E-F). Despite a large panel of up-regulated cytokine mRNA in MC^TH^, we did not find evidence that MC^TH^ were able to produce IL12B, coding for IL-12 p40 subunit of key polarizing IL-12 and IL-23 cytokines (Figure 1F). Both IL-12 and IL-23 were also not detected in MC^TH^ supernatant (Figure S1G).

To further extend the characterization of the impact of activated Teff cells on MCs, we analyzed key mast cells functions (*e.g*. degranulation and eicosanoid production) following interaction with activated CD4^+^ T cells. Anti-DNP IgE sensitized MCs were co-cultured with CD4^+^ T cells with or without anti-CD3/CD28 coated beads for 48 hours and next challenged with DNP-BSA for 30 minutes (Figure 2A). Whilst FcεRI gene expression was not impacted by activated CD4^+^ T cells (table S2), MCs showed a decreased FcεRI activation threshold and an increased degranulation capacity (Figure 2A). The analysis of 27 polyunsaturated fatty acid metabolites by liquid chromatography coupled to tandem mass spectrometry (Le Faouder et al., 2013) showed that when MCs were co-cultured with activated CD4^+^ T cells only COX-dependent eicosanoids were increased (Thromboxane B_2_, Prostaglandin E_2_ (PGE_2_), PGD_2_, 8-isoPGA_2_, 15d-PGJ_2_) (Figure 2B).

**Figure 2.**
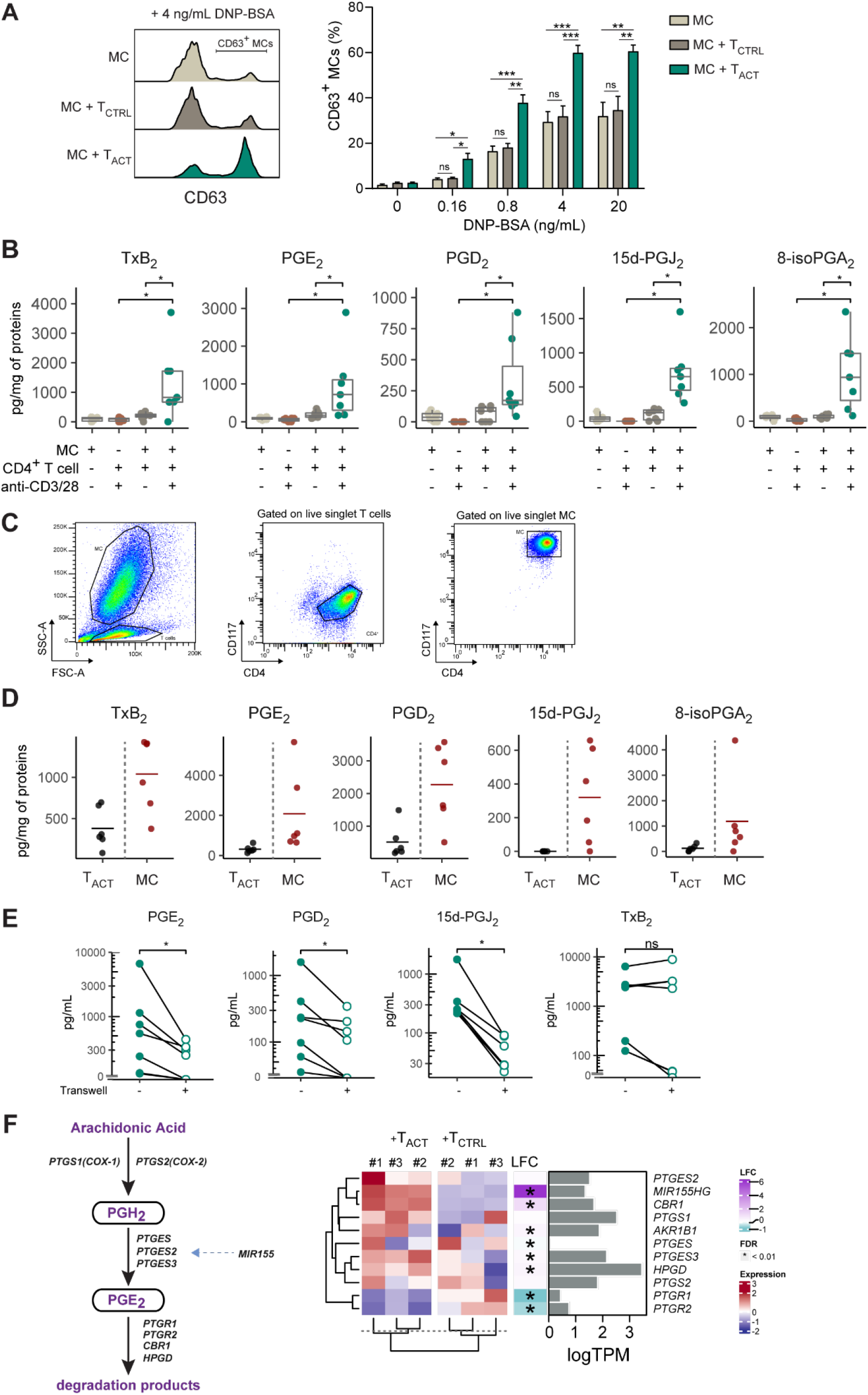
Activated CD4^+^ T cells potentiate MC degranulation and induce eicosanoid production. (A) anti-DNP IgE sensitized MCs were cocultured with T_ACT_ or T_CTRL_ for 48 hours and next challenged with DNP-BSA for 30 min. Degranulation was measured by CD63 exposure on the MC surface by flow cytometry. Representative histograms (left panel) and pooled data (mean ± SEM, n=8) (right panel) from 3 independent experiments. Two-way ANOVA with Tukey’s multiple comparisons test. (B-D) MCs were cocultured with T_ACT_ or T_CTRL_ for 48 hours. Eicosanoids were quantified in the cell pellets by LC-MS/MS. Data are presented as box and whiskers plot (Tukey style), each point represents a MC/T cell pair (n=7 from 3 independent experiments). Friedman test and pairwise comparisons using paired Wilcoxon signed-rank test. (B). MCs and CD4^+^ T cells were FACS-sorted from +T_ACT_ condition (FACS gating strategy, C) and eicosanoids were quantified in the cell pellets by LC-MS/MS. Bars represent the mean (n=6), each point represents a MC/T cell pair (D). (E) MCs were cocultured with T_ACT_ separated or not with 0.4 µm transwell inserts for 48 hours. Eicosanoids were quantified in the cell supernatant by LC-MS/MS. Paired Wilcoxon signed-rank test. *p < 0.05, **p < 0.01, ***p < 0.001, ****p < 0.0001, ns not significant. (F) Scheme of PGE_2_ metabolism and related genes (dashed arrows indicate indirect positive action) and clustered heatmap showing the relative expression level of each MC gene between MC +T_ACT_ vs MC +T_CTRL_ conditions. Each gene expression level in +T_ACT_ condition (as depicted by transcript per kilobase million, TPM) and its log_2_-fold-change expression and associated FDR (stars indicate FDR<0.01) are also shown.

Because both effector CD4^+^ T cells and MCs are able to produce eicosanoids, we quantified eicosanoids in each cell type isolated by FACS cell sorting based on SSC^low^/FSC^low^/CD117^-^ /CD4^+^ and SSC^high^/FSC^high^/CD117^+^/CD4^+^ profiles, respectively (Figure 2C-D). Eicosanoids increased in coculture condition were mainly detected in MCs compared to activated CD4^+^ T cells; only very low amounts of some lipid mediators (e.g., TxB_2_ and PGD_2_) were found in T cells (Figure 2D). These results indicated that activated effector CD4^+^ T cells triggered COX-dependent eicosanoids production in MCs. Similar co-culture experiments performed with Transwell plates showed that PGE_2_, PGD_2_, 15d-PGJ_2_ production required direct contact between MCs and CD4^+^ T cells whereas TxB_2_ production was contact-independent (Figure 2E).

The inducible key rate-limiting enzyme in prostanoid synthesis, COX-2, being not upregulated in MC^TH^ (Table S2), we analyzed the expression levels of genes coding for enzymes directly involved in PGE_2_ metabolism as well as MIR155, a regulator of this pathway (Kim et al., 2021). Our analysis indicated that PGE_2_ production in MC^TH^ may be dependent on both the up regulation of prostaglandin E synthases (PTGES2/3) and a downregulation of catabolic enzymes HPGR1 and HPGR2 (Figure 2F). In agreement, *MIR155* was also upregulated suggesting its role as indirect positive regulator of prostaglandin E synthase (*PTGES2/3*) expression.

Taken together these results show that activated Th cells induce a specialized MC population with a specific genetic program resulting in acquired functionalities notably the production of cytokines and eicosanoid mediators.

### MC^TH^ drive Th cells toward IL-17 production in an eicosanoid-dependent manner

Because MC^TH^ exhibit an increased capacity to interact with T cells and they produce eicosanoids, we investigated the impact of MC^TH^ on Teff cell responses. Even if PGE_2_ is the main eicosanoid described to impact Th cell responses (Tsuge et al., 2019), contradictory mechanisms were reported indicating that, eicosanoids impact on Th cells is fine-tuned and strongly depend on their development state, other cells and cues of the environment (Dejani et al., 2018; Kofler et al., 2014; Lee et al., 2019; Yao et al., 2009). To assess the influence of MC derived-eicosanoids on CD4^+^ T cell responses, we used indomethacin, an irreversible cyclooxygenase inhibitor. We first validated that addition of indomethacin (100 µmol/L), at the beginning of MC/Th cell cocultures nullified PGE_2_, PGD_2_ and 15 dPGJ_2_ production (figure S2A).

We monitored the proliferation and CD25 expression kinetics of effector/memory CD4^+^ T cells activated by anti-CD3/CD28 coated beads when incubated or not with MCs in the presence or the absence of indometacin. MCs greatly enhanced CD4^+^ T cell proliferation via eicosanoids (Figure 3A). By contrast, CD25 up regulation was independent on eicosanoids and its expression on CD4^+^ T cell surface was slightly increased in presence of MCs at the beginning of the coculture but reached similar levels after the fifth day (Figure 3B and Figure S2B). Of note, indomethacin did not impact T cells proliferation or CD25 expression in absence of MCs (Figure 3A-B). We next analyzed CD4^+^ T cell subset distribution based on their cytokine production profiles at day 6 of coculture corresponding to T cell proliferation and activation plateau. Intracellular cytokines were analyzed after PMA/ionomycine restimulation. Th cell subsets were defined according to the panel of cytokines produced: Th1 as IFN-γ^+^/IL-17^-^/IL-22^-^, Th2 as IFN-γ^-^/IL4^+^, Th17 as IFN-γ^-^/IL-17^+,^ Th22 as IFN-γ^-^/IL-17^-^/IL-22^+^ and Th1.17 as IFN-γ^+^/IL-17^+^. In parallel experiments, Treg were gated as FOXP3^+^, LAP^+^, CD25^+^ and CD127^low^, without restimulation (Figure S2C). Because Ag-experienced CD4^+^ T cells isolated from human blood can have very different priming history that might influence next the interplay with MC in the coculture, we analyzed more than 70 MC-CD4^+^ Teff cell pairs from different donors. To globally analyze the impact of MCs on the Th subset distribution, we first performed a PCA (Figure S2D) on the percentage of each T cell subset. This unsupervised analysis revealed that MCs shifted the distribution of the T cell subsets toward Th17 and Th1.17. Addition of indometacin in the MC/Th cell coculture shifted the T cell subset distribution mainly towards the Th2 loading. Indometacin showed no impact when added in Th cells activated in the absence of MCs (Figure S2D).

**Figure 3.**
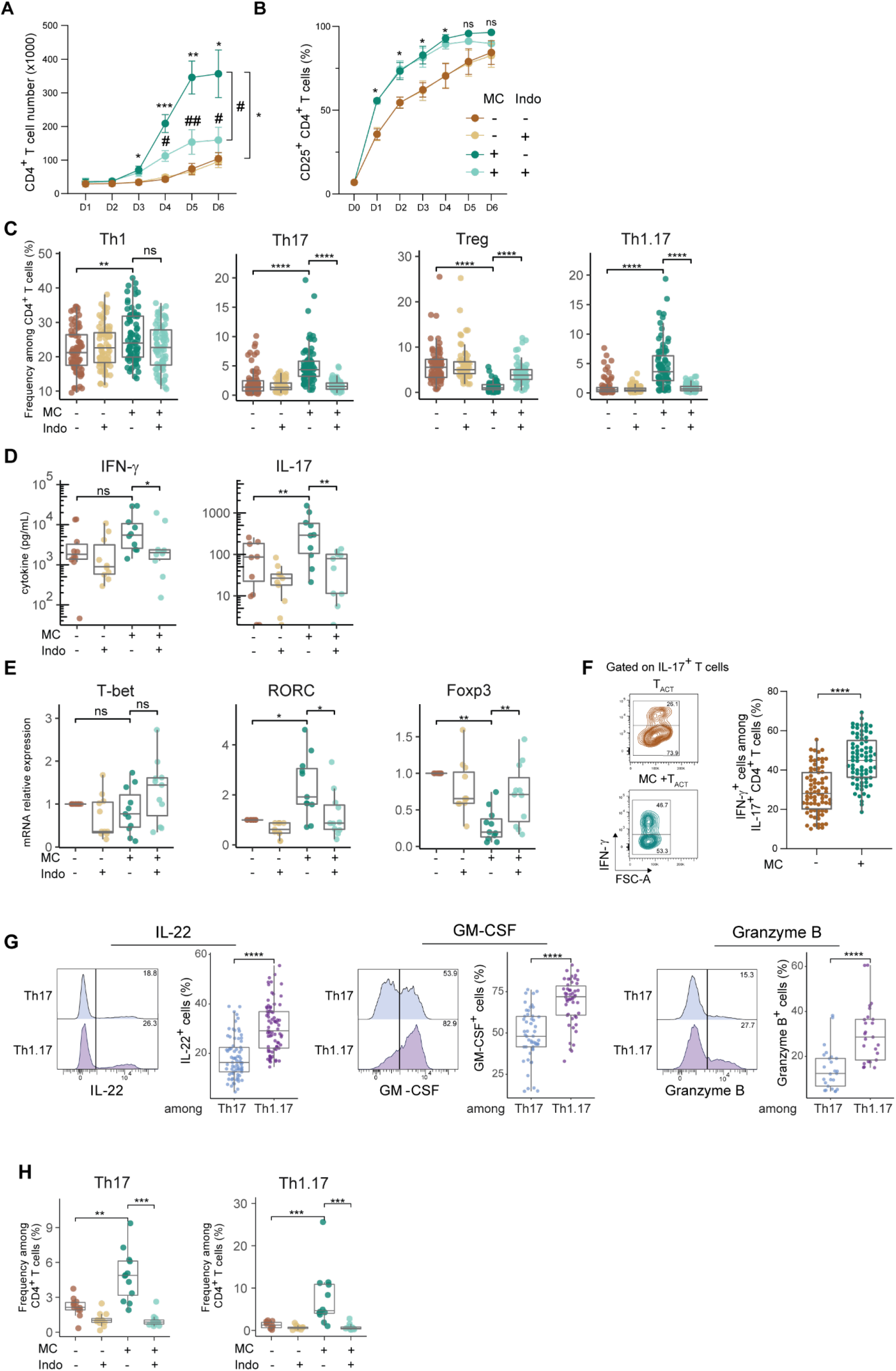
Helped hMCs drive Th cell toward IL-17 production in a COX-2 dependent manner. Effector/memory CD4^+^ T cells were cocultured with MCs in presence of anti-CD3/CD28 coated beads for 6 days with or without indometacin. (A-B) Kinetics of CD4^+^ T cell numbers (A) and CD25 expression in CD4^+^ T cells (B). Pooled data from 9 (A) and 6 (B) independent experiments (mean ± SEM, two-way ANOVA with Dunnet post hoc test, compared conditions: MC +T_ACT_ vs T_ACT_ (*) and MC+T_ACT_ vs MC+T_ACT_ +indo (#)). (C) Th cell subset analysis at day 6 of coculture. Each subset frequencies are presented as box and whiskers plot (Tukey style), each point represents a MC/T cell pair. Pooled data (n=77) from 31 independent experiments, Friedman test and pairwise comparisons using Dunn’s test. (F) (D-E) After 6 days coculture, CD4^+^ T cells were FASC-sorted. Cytokine amounts were measured after restimulation with PMA and ionomycin (n=9 from 3 independent experiments) (D). Transcription factors expression was assessed by RT-qPCR (n=12 from 8 independents experiments) (E). Data are presented as box and whiskers plot (Tukey style), each point represents a MC/T cell pair. Friedman test and pairwise comparisons using paired Wilcoxon signed-rank test. (F) Frequency of IFN-g^+^ cells among IL-17^+^ CD4^+^ T cells. Representative dot plots and pooled data (n=77) from 31 independent experiments. Data are presented as box and whiskers plot (Tukey style), each point represents a MC/T cell pair, paired Wilcoxon signed-rank test. (G) Flow cytometry analysis of GM-CSF, IL-22 and Granzyme B expression by Th17 or Th1.17 cell subsets at day 6 of coculture with MCs. Representative histograms and pooled data (n=25 to 77 from 5 to 31 independents experiments). Data are presented as box and whiskers plot (Tukey style), each point represents a MC/T cell pair, paired Wilcoxon signed-rank test. (H) Flow cytometry analysis of Th17 and Th1.17 subset frequencies at day 12 of coculture. Data are presented as box and whiskers plot (Tukey style), each point represents a MC/T cell pair from 8 independents experiments, Friedman test and pairwise comparisons using paired Wilcoxon signed-rank test. *p < 0.05, **p < 0.01, ***p < 0.001, ****p < 0.0001, ns not significant. See also Figure S2.

Analysis of each CD4^+^ T cell subset frequency (Figure 3C and S2E) showed that Th1, Th2, Th17, Th1.17 subsets were increased in the presence of MCs while Treg and Th22 frequencies were decreased. Of note, we observed that Th17 and Th1.17 profiles were the most impacted by MCs. Indometacin completely abolished the effect of MCs on Th17 and Th1.17 frequencies whereas it exhibited a mild effect on Th1 cells. On the contrary, indometacin treatment led to an increase in Th2 cells and dampened the negative impact of MCs on Treg frequency. The reduction of the Th22 cell frequency observed in the presence of MCs was reinforced by indometacin (Figure 3C and S2E). Flow cytometry analysis was confirmed by measuring the cytokine amounts produced by CD4^+^ cells isolated and then restimulated with PMA/ionomycine for 6 hours. The quantities of IFN-γ, IL-4 and IL-17 released by activated CD4^+^ T cells were increased in the presence of MCs. The release of IFN-γ, IL-17 and IL-22 was decreased by indometacin while that of IL-4 was increased (Figure 3D and S2F).

Expression of Th master transcription factors quantified by RTqPCR on FACS isolated CD4^+^ T cells showed that whilst RORC expression was increased in the presence of MCs, both FOXP3 and AhR were decreased. The effects of MCs on the expression of RORC and FOXP3 were reversed by indomethacin. As observed for Th2 cytokines, GATA3 expression was enhanced in the presence of MCs plus indometacin (Figure 3E and S2G). These results clearly indicate that MC^TH^-derived eicosanoids promote the emergence of specific T cell subsets. A more detailed analysis of IL-17^+^ CD4^+^ T cell population showed that the fraction of Th1.17 (i.e. IFN-γ^+^) among IL-17^+^ cells doubled in the presence of MCs (Figure 3F).

To further characterize the Th1.17 cells induced by MCs, we measured the expression of IL-22, GM-CSF and Granzyme B previously identified as strongly expressed in pathogenic Th1.17 as compared to non-pathogenic Th17 cells (Lee et al., 2012). Th1.17 cells induced in the presence of MC expressed more frequently IL-22, GM-CSF and Granzyme B (Figure 3G) than their Th17 counterpart. Finally, we investigate the stability of the IL-17^+^ Th cells generated in the presence of MCs. Following 12 days of co-culture between MCs and CD4+ T cells, the analysis of intracellular cytokines showed that Th17 and Th1.17 profiles were maintained in an eicosanoid dependent manner (Figure 3H and S2H). These results suggest that MC^TH^ derived eicosanoids have a long imprinting on the profiles of Teff cell responses.

Taken together, these data suggest that MC^TH^ interplay with activated Th cells leads to eicosanoid production by MCs that in turn shapes Th cell subset by on the one hand promoting IL-17 producing subsets and notably Th1.17 cell subsets and on the other hand dampening Treg and Th2 subsets.

### Th1.17 cells emerge from a distinct CCR6^+^ CD4^+^ T cell population in presence of _MC_TH

To identify the origin of IL-17-producing T cells induced by MCs, we first analyzed whether MCs conferred to IL-17^+^ cell subsets a proliferative benefit. CTV labelled CD4^+^ T cells were cocultured for 4 days with MCs in presence of anti-CD3/CD28 coated beads. The round number of divisions was analyzed in each Th cell profile (Figure S3A-B). Whilst MCs supported the proliferation of all Th cells (in accordance with data in Figure 3A), the division profiles obtained for each Th cell subset were similar, indicating that in the presence of MCs, the different Th cell subsets divided at a similar pace (Figure S3A-B).

Because Treg cells can differentiate into Th17 cells (Koenen et al., 2008; Ueno et al., 2013), we assessed whether Treg may turn into IL-17 producing cells in our experimental setting. For this purpose, Treg cells were sorted, fluorescently tagged and reincorporated with their Tconv counterparts. Following interaction with MC^TH^, some Th17 and Th1.17 emerged from Treg cells indicating that MCs can induce the differentiation of IL-17 producing cells from Treg cells (Figure S3C-D). Nevertheless, these IL-17^+^ cells derived from Treg cells accounted only for a minor percentage of IL-17^+^ cells in the whole CD4^+^ T cell population (< 3%) indicating that the vast majority of IL-17^+^ Th cells induced by MCs derived rather from Tconv cells than Treg cells (figure S3E).

These results prompted us to investigate Tconv subpopulation from which originate IL-17^+^ cells. The main Th subsets were sorted according to the expression of chemokine receptors as described in the literature (CXCR3^-^CCR4^-^CCR6^-^CCR10^-^ as Th0, CXCR3^+^CCR4^-^CCR6^-^ CCR10^-^ as Th1, CXCR3^-^CCR4^+^CCR6^-^CCR10^-^ as Th2, CXCR3^-^CCR4^+^CCR6^+^CCR10^-^ as Th17 and CXCR3^-^CCR4^+^CCR6^+^CCR10^+^ as Th22) and cocultured with anti-CD3/CD28 beads in presence of MCs for 6 days (Figure 4A). IL-17^+^/IFN-γ^-^ cells (Th17) were detected only in experiments with CXCR3^-^CCR4^+^CCR6^+^CCR10^-^ and CXCR3^-^CCR4^+^CCR6^+^CCR10^+^ cell populations and MC^TH^ increased their frequencies in an eicosanoid dependent manner (Figure 4B-C). Furthermore, MC^TH^-derived eicosanoids induced IL-17^+^/IFN-γ^+^ cells (Th1.17) only from the CXCR3^-^CCR4^+^CCR6^+^CCR10^-^ cell population (Figures 4B-D).

**Figure 4.**
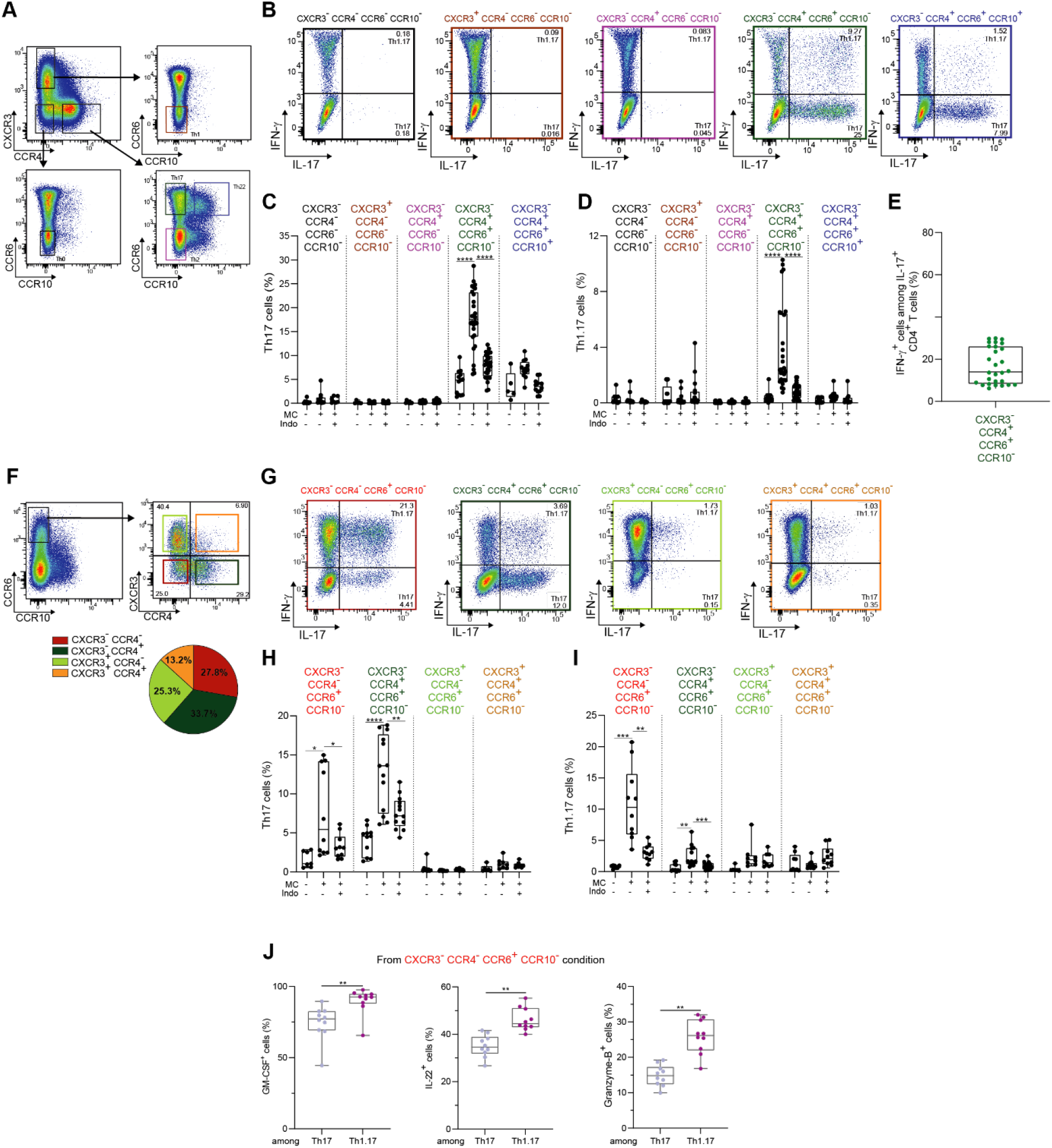
Origin of IL-17^+^ Th cells during MC/Th cell interplay. Th cell subsets were FACS-sorted according to indicated chemokine receptor expression. (A) cell sorting strategy (B-E) Sorted Th cell subsets were cocultured with MCs ± indometacin for 6 days and IFN-γ and IL-17 expression was assessed by flow cytometry following PMA/ionomycin restimulation. Representative dot plots of MCs + T_ACT_ condition (B) and pooled data of Th17 (C) and Th1.17 (D) frequencies in indicated conditions. Data are presented as box and whiskers plot (min to max), each point represents a MC/T cell pair (n=11 to 28). Frequency of IFN-g^+^ cells among IL-17^+^ T cells in cocultures with CXCR3^-^CCR4^+^CCR6^+^CCR10^-^ T cells (E). Mean ± SEM each point represents a MC/T cell pair (n=11 to 28). (F) Cell sorting strategy and frequency of CXCR3/CCR4 subsets among CCR6^+^ CCR10^-^ CD4^+^ T cells (n=10) (G-J) Sorted Th cell subsets were cocultured with MCs ± indometacin for 6 days and IFN-γ and IL-17 expression was assessed by flow cytometry following PMA/ionomycin restimulation. Representative dot plots of MCs + T_ACT_ condition (G) and pooled data of Th17 (H) and Th1.17 (I) frequencies in indicated conditions. Data are presented as box and whiskers plot (min to max), each point represents a MC/T cell pair (n=8 to 13). GM-CSF, IL-22, Granzyme B expression among Th17 and Th1.17 cells from coculture with CXCR3^-^CCR4^-^CCR6^+^CCR10^-^ T cells and MCs (J). Paired Wilcoxon signed-rank test. *p < 0.05, **p < 0.01, ***p < 0.001, ****p < 0.0001. See also Figure S3

Nevertheless, in CXCR3^-^CCR4^+^CCR6^+^CCR10^-^ T cells and MC^TH^ coculture, we observed that IFN-γ^+^ cells accounted for only 20% of IL-17^+^ cells which is lower compared to the ∼50% observed by using the whole CD4^+^ T cell population (Figure 3I). This suggested that another population contributes to the emergence of Th1.17 cells (Figure 4E). Because transcriptionally different CCR6^+^ Th17 subsets have been previously reported (Wacleche et al., 2016), coculture experiments were carried out on four CCR6+CCR10-subsets defined relative to their expression of CCR4 and CXCR3 (Figure 4F). Following 6 days of coculture with MC^TH^, Th17 cells emerged from CXCR3^-^CCR4^-^CCR6^+^CCR10^-^ and CXCR3^-^CCR4^+^CCR6^+^CCR10^-^ cells and most of Th1.17 cells derived from the CXCR3^-^CCR4^-^CCR6^+^CCR10^-^ subpopulation. All the effects of MC on Th17 subsets were dependent on eicosanoid production (Figure 4G-I). Moreover, the Th1.17 cells that emerged from the CXCR3^-^CCR4^-^CCR6^+^CCR10^-^ subpopulation expressed the pathogenicity markers IL-22, GM-CSF and Granzyme B (Figure 4J).

Collectively, these results show that MCs increase the frequency of IL-17-producing (CXCR3^-^ CCR4^+^CCR6^+^CCR10^-^) Th17 cells and allow the emergence of Th1.17 cells with a pathogenic phenotype from a specific CD4^+^ T cell subset (CXCR3^-^CCR4^-^CCR6^+^CCR10^-^).

### MC^TH^ derived PGE_2_ combined to IL-1β promote Th1.17 cells

To identify which COX-2 dependent eicosanoids promote the induction of IL-17^+^ CD4^+^ T cells by MC^TH^, we complemented indometacin-treated MC/Th cell coculture with increasing concentration of PGE_2_, PGD_2_, 15d-PGJ_2_. When T cells were stimulated in presence of MC^TH^ and indomethacin, the addition of PGE_2_ restored the frequencies of Th17, Th1.17, Th22, Th2 and Treg in a dose dependent manner while Th1 frequencies remained unchanged (Figure 5A-B and S4A). In contrast, neither PGD_2_ nor 15d-PGJ_2_ addition showed any effect on the Th cell subset frequencies (Figure S4B-C). Based on RNAseq dataset showing that activated CD4^+^ Teff cells express the PGE_2_ receptors subclasses EP2, EP3 and EP4 (LaMere et al., 2017), we investigated the role of each PGE_2_ receptor subtype by adding specific agonists in coculture experiments (butaprost for EP2, sulprostone for EP3 and L-902,688 for EP4 (Markovic et al., 2017)). Butaprost and L-902,688 with a lower efficacy but not sulprostone reproduced the PGE_2_ effects on subsets Th17, Th1.17 and Treg frequencies (Figure 5C-D) and S4D). Collectively, these results indicated that the effects of MC^TH^ eicosanoids on Th cell subset distribution is orchestrated by the action of PGE_2_ mainly through EP2 signaling.

**Figure 5.**
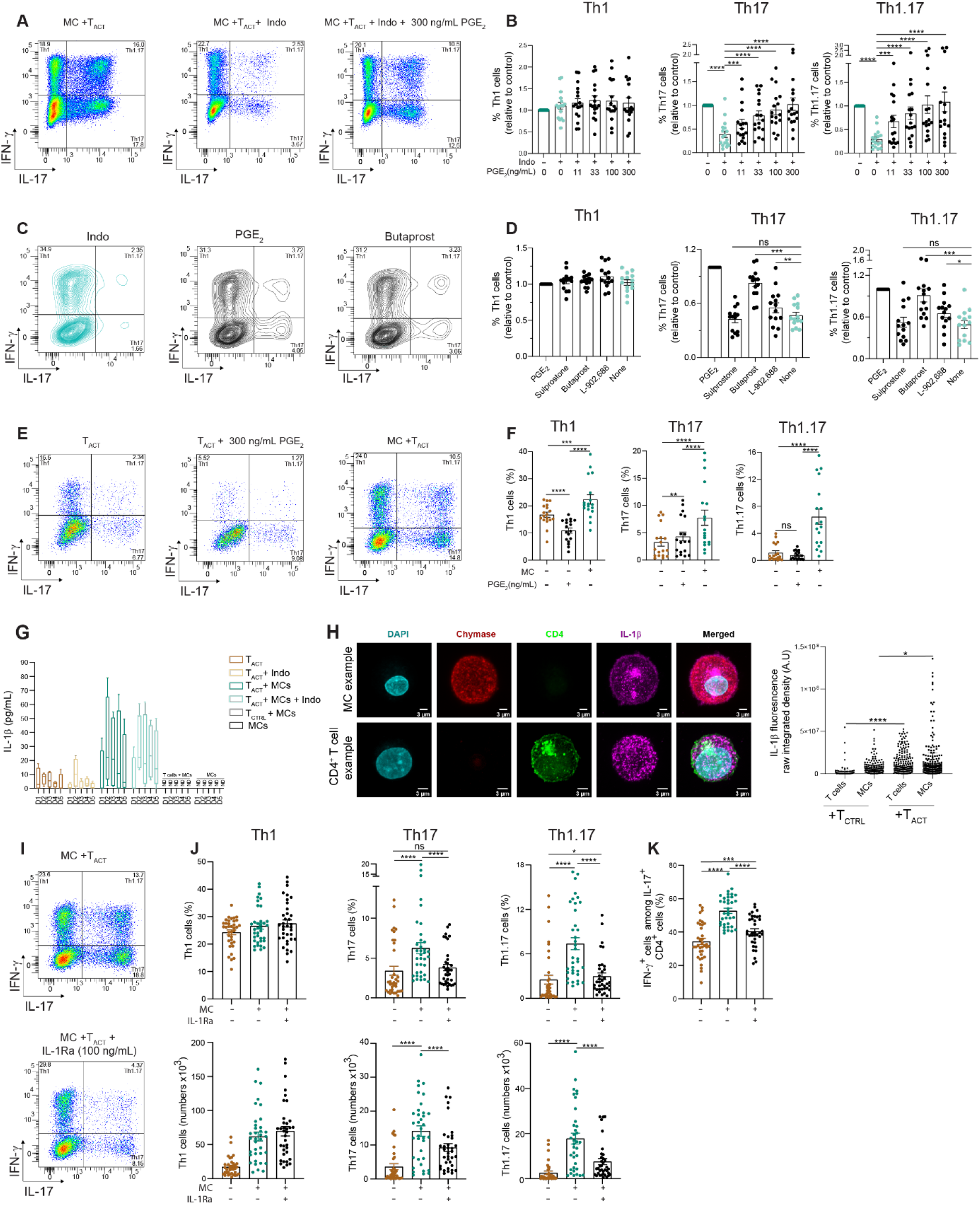
PGE_2_ and IL-1β are required for the induction of IL-17 producing Th cell subsets. Effector/memory CD4^+^ T cells were cultured with or without MCs in the presence of anti-CD3/CD28 coated beads for 6 days. (A-B) Indometacin-treated cocultures were supplemented with increasing concentrations of PGE_2_ and Th cell subsets frequencies were analyzed by flow cytometry at day 6. Dot plots of one representative experiment (A), Th cell subset frequencies are presented as fold change over the control condition (MC +T_ACT_), mean ± SEM, each point represents an experiment (n=17 from 13 independent experiments) (B). (C-D) Indometacin-treated cocultures were supplemented with PGE_2_ receptor agonists or 300 ng/mL of PGE_2_. Th cell subsets frequencies were analyzed by flow cytometry at day 6. Dot plots of one representative experiment (C). Th cell subset frequencies are presented as fold change over the control condition (T_ACT_ + MC with indometacin and 300 ng/mL PGE_2_), mean ± SEM, each point represents an experiment (n=14 from 4 independent experiments) (D). (E-F) Th cell subsets frequencies at day 6 in PGE_2_ +T_ACT_ as compared to MC +T_ACT_ condition. Dot plots of one representative experiment (E). Mean ± SEM of Th cell frequencies (n=18 from 8 independent experiments), each point represents one experiment (F). (G) Kinetics of secreted IL-1β measured in supernatants of indicated conditions. Mean ± SEM from 4 independent experiments. (H) Confocal microscopy images of IL-1b^+^ MCs and IL-1b^+^ CD4^+^ T cells after 2 days of coculture showing staining (average intensity projections) for Chymase (red), IL-1β (magenta), CD4 (green) and DAPI (cyan), and quantification of IL-1β fluorescence integrated density in MCs and CD4^+^ T cells (n=3 independent experiments). Kruskal Wallis and pairwise comparisons using Dunn’s test (I-K) Th cell subsets frequencies at day 6 in IL1Ra-treated cocultures. Dot plots of one representative experiment (I). Frequencies and numbers of Th cell subsets (J). Frequency of IFN-g^+^ cells among IL-17^+^ CD4^+^ T cells (K). Mean ± SEM, each point represents a MC/T cell pair (n=36 from 8 independents experiments) Friedman test and pairwise comparisons using paired Wilcoxon signed-rank test (except in H). *p < 0.05, **p < 0.01, ***p < 0.001, ****p < 0.0001. See also Figure S4.

To assess whether the impact of MCs on Th17 and Th1.17 subsets could be mimicked by PGE_2_, we carried out similar experiments on CD4^+^ Th cells activated without MC in the presence of 300 ng/mL PGE_2_. PGE_2_ alone increased the percentages of Th17 cells but to a lesser extent to that observed in the presence of MC (Figure 5E-F). PGE_2_ alone did not induce the emergence of Th1.17 cells that required the presence of MCs (Figure 5E-F). We then hypothesized that another mediator from MC^TH^ is required to generate Th1.17. Because *IL1B* gene is expressed by MC^TH^ and that IL-1β is known to participate to the induction of IL-17^+^ Th subsets (Acosta-Rodriguez et al., 2007a), we quantified the release of IL-1β in our experimental setting. IL-1β was produced only when activated CD4^+^ T cells were cocultured with MCs independently of eicosanoid production (Figure 5G). Moreover, IL-1β was produced by both cell types at 48 hours of coculture (Figure 5H). IL-1 receptor antagonist (IL-1Ra) was next used to examine the impact of IL-1β on Th17 and Th1.17 cell subsets. Addition of IL-1Ra in the cocultures reduced the frequencies and the numbers of Th17 and Th1.17 cell subsets without affecting Th1 cell subsets (Figure 5I-J). In addition, the frequency of Th1.17 among IL-17-producing Th cells was reduced by IL-1Ra treatment indicating that IL-1β is a key factor in the induction of Th1.17 cell subsets by MCs (Fig 5K).

Together, these results identify PGE_2_ and IL-1β as two key factors of the MC-CD4^+^ T cell interplay that contribute to the emergence of Th17 and Th1.17 cell subsets.

### MCs exhibit a COX-2^+^/IL-1β^+^ profile in colonic mucosa of patients with Crohn disease

Since MC/Teff cell interplay allows the emergence of Th1.17 cells that are involved in immune-mediated inflammatory disease (IMID) including auto-immune diseases, we used Open Targets Platform (an open access integrated multiomics data resource that allows to link genes/proteins to a disease of interest, https://platform-docs.opentargets.org/) (Ochoa et al., 2020) to determine whether genes upregulated in MC^TH^ overlapped significantly with gene linked to IMID. This analysis indicated that multiple IMID-associated genes overlapped significantly with MC^TH^ upregulated genes, such as ulcerative colitis, Crohn disease, vitiligo, psoriasis, multiple sclerosis, lupus, and rheumatoid arthritis (Figure 6A) suggesting a possible implication of MC/Teff cell interplay in these pathologies. We focused on IBD and found that 47 genes overlapped between upregulated genes in MC^TH^ and a curated GWAS-prioritized gene list (Mukhopadhyay et al., 2019) (Figure 6B and table S4). Among them we found upregulated genes (table S2) and key transcription factors (table S3) identified in MC^TH^

**Figure 6.**
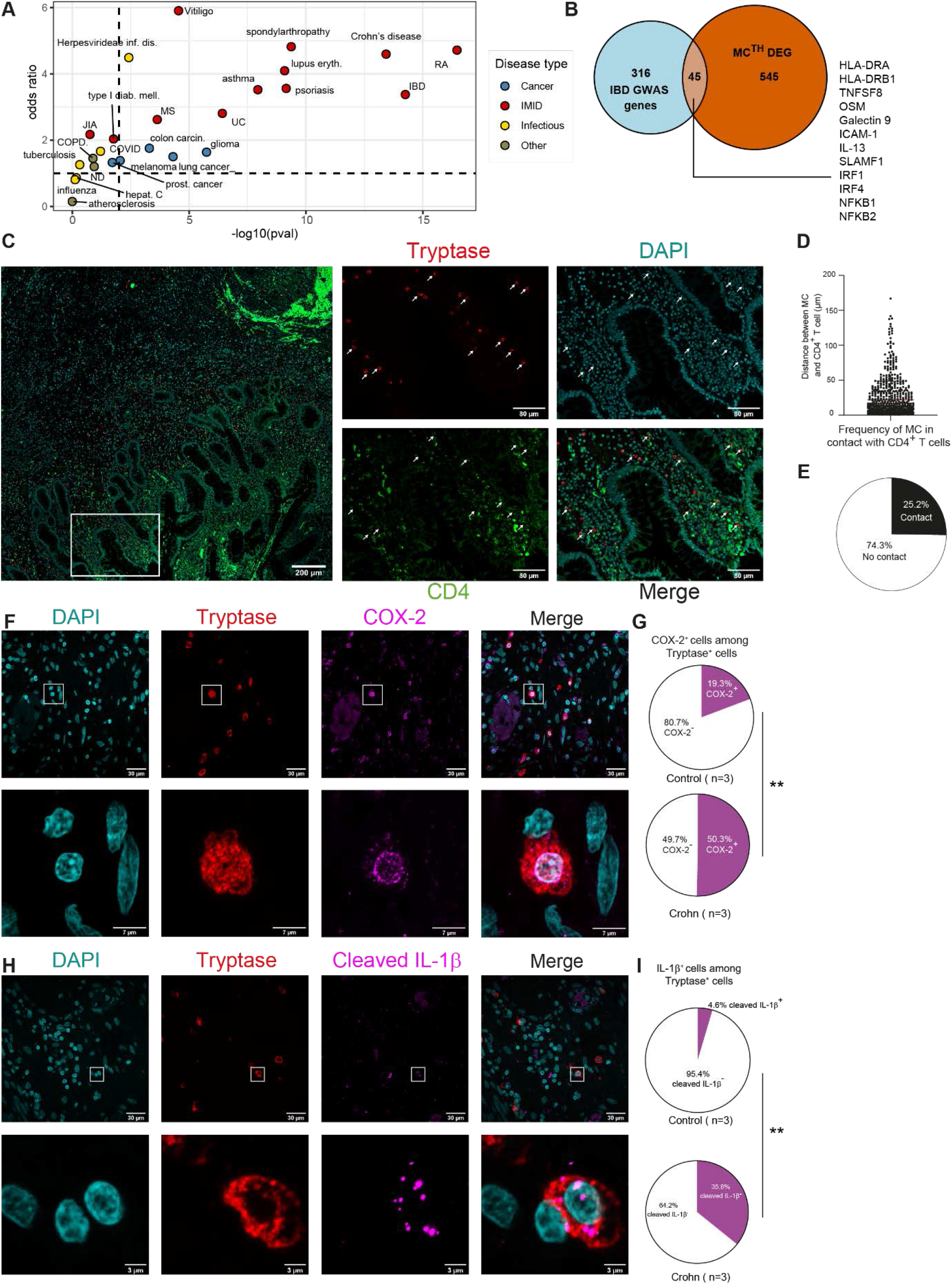
Crohn’s disease-associated MCs exhibit a COX-2^+^/IL-1β^+^ profile. (A-B) Overlap between upregulated genes in MC^TH^ and genes associated with IMIDs or other diseases (retrieved via the Open Target Platform, JIA, juvenile idiopathic arthritis; MS, multiple sclerosis; ND, neurodegenerative diseases; RA, rheumatoid arthritis; UC, ulcerative colitis). Overlap (tested using geneOverlap R Package) for the different pathologies tested was presented by plotting odd ratio (which gives the strength of the association) vs P-value (contingency tables) (A). Overlap between upregulated genes in MC^TH^ and prioritized gene candidates identified in IBD GWAS is shown as proportional Ven’s diagram. Odd ratio= 6.29 and pval. = 6.9 10^-20^. Shown are some overlapping genes, see also Table S4 (B). (C-E) MC and CD4^+^ T cell distribution in colonic mucosa from patients with Crohn’s disease. Paraffin embedded tissues were stained for CD4 and tryptase markers. Representative staining image presented in maximal fluorescence intensity projection, box indicates the region depicted in the right panels, scale bars, scale bars 200 µm (left panel) and 80 µm (right panel) (C). Measure of the distances between randomly selected MCs and their proximal CD4^+^ T cell (n=636 from 6 independent experiments) (D). Percentage of MCs in direct contact with CD4^+^ T cells. MCs of the imaged area were analyzed in 6 independents experiments (E). (F-G) Colonic mucosa from Crohn’s disease or control patients were stained for CD4 (green), Tryptase (red), COX-2 (magenta) and DAPI (cyan). Representative images from patient with Crohn’s disease, boxes indicate the region depicted in the lower panels (F). Quantification of COX-2^+^ MCs (n=3, G). (H-I) Colonic mucosa from Crohn’s disease or control patients were stained for CD4 (green), Tryptase (red), cleaved IL-1β (magenta) and DAPI (cyan). Representative images from patient with Crohn’s disease, boxes indicate the region depicted in the lower panels (H). Quantification of COX-2^+^ MCs (n=3, I). Unpaired *t*-test, ** p<0.01. See also Table S4.

Accordingly, we investigated MC/CD4^+^ T cell interplay in biopsies from patients with Crohn’s disease. High resolution tile-scan imaging revealed that MCs (tryptase^+^) were often at the vicinity of CD4^+^ T cells (Figure 6C). Quantitative analysis of more than 600 MCs indicated that the mean-distance between MCs and T cells was ∼22 µm, a distance compatible for paracrine interactions ((Müller et al., 2012), Figure 6 D). Around 25% of MCs were in direct contact with CD4^+^ T cells (Figure 6E). The histological analysis of the expression of COX-2 and the mature form of IL-1β (cleaved IL-1β) showed that the frequency of MCs expressing COX-2 and mature IL-1β was dramatically increased within biopsies from Crohn’s patients as compared to control samples (50.3% versus 19.3% and 35.8% versus 4.6% respectively) (Figure 6F-I).

Taken together, these observations suggest that pro-inflammatory MCs, characterized by the expression of COX-2 and IL-1β, that support IL-17 production by CD4^+^ T cells are hugely increased in Crohn’s disease.

### Late-stage colitis severity is reduced in KIT-deficient mice

The impact of MC/Th cell interplay was assessed in KIT-deficient mice (Kit^W-sh/W-sh^) (Saleh et al., 2019) in the DSS-induced colitis model involving colitogenic IL-17-producing CD4^+^ T cells. Colitis was induced by adding dextran sulfate sodium (DSS, 3%) into the drinking water for 5 days and then followed by 5 days of drinking water. In the first 6 days of colitis (acute phase) Kit^W-sh/W-sh^ and WT littermate mice (KIT^+/+^) lost weight similarly. From the 7^th^ day of the colitis (2 days after removing DSS from drinking water), loss of body weight was restoring in Kit^W-sh/W-sh^ MC-deficient mice whereas control WT mice continued to lose weight (Figure 7A). Inflammation-induced colonic tissue damage was similar between the two strains of mice at day 5 of the disease and was reduced only in Kit^W-sh/W-sh^ at day10 (Figure 7B-F). Of note, no differences were observed between males and females irrespective of their genotype (Figure 7 B-E). We next investigated inflammatory cytokine and chemokine expression in the colon at day 5 and day 10. We observed a decrease of *Ifng, Cxcl9* and *Cxcl10* gene expression level in Kit^W-sh/W-sh^ at day 5 (Figure 7G-H). In agreement with a reduced colitis severity at day 10 of the disease in Kit^W-sh/W-sh^ mice, gene expression levels of all the inflammatory cytokines and chemokines including *Il1b, Il6, Il17a, Il22, Ifng, Ccl1, Ccl5, Cxcl2, Cxcl9, Cxcl10* were lower in Kit^W-sh/W-sh^ as compared to their WT littermates (Figure 7G-H). As we showed that MC-derived eicosanoids were involved in promoting IL-17 production by Th cells, we analyzed the expression of key enzymes in eicosanoid metabolism. Although lipoxygenase expression (i.e. *Alox5, Alox12* and *Alox15*) was similar in both strain, COX-2 expression (i.e. *Ptgs2*) was lower in KIT-deficient mice (Figure 7I). In line with a role of MC in the induction/maintenance of colitogenic T lymphocytes, mucosal T cell density was strongly reduced in the colon of KIT-deficient mice at day 10 (Figure 7J-L). Within *lamina propria* of WT mice, the majority of the MCs were within a distance of 50 µm of T cells and 20 % of mucosal MC were in direct contact with T cells suggesting possible paracrine, juxtacrine or synapse-mediated cell-cell communication (Figure 7M-N). Collectively, these results indicated that, in absence of MCs, mice exhibited a mitigated pathology notably in the late phase of the disease.

**Figure 7.**
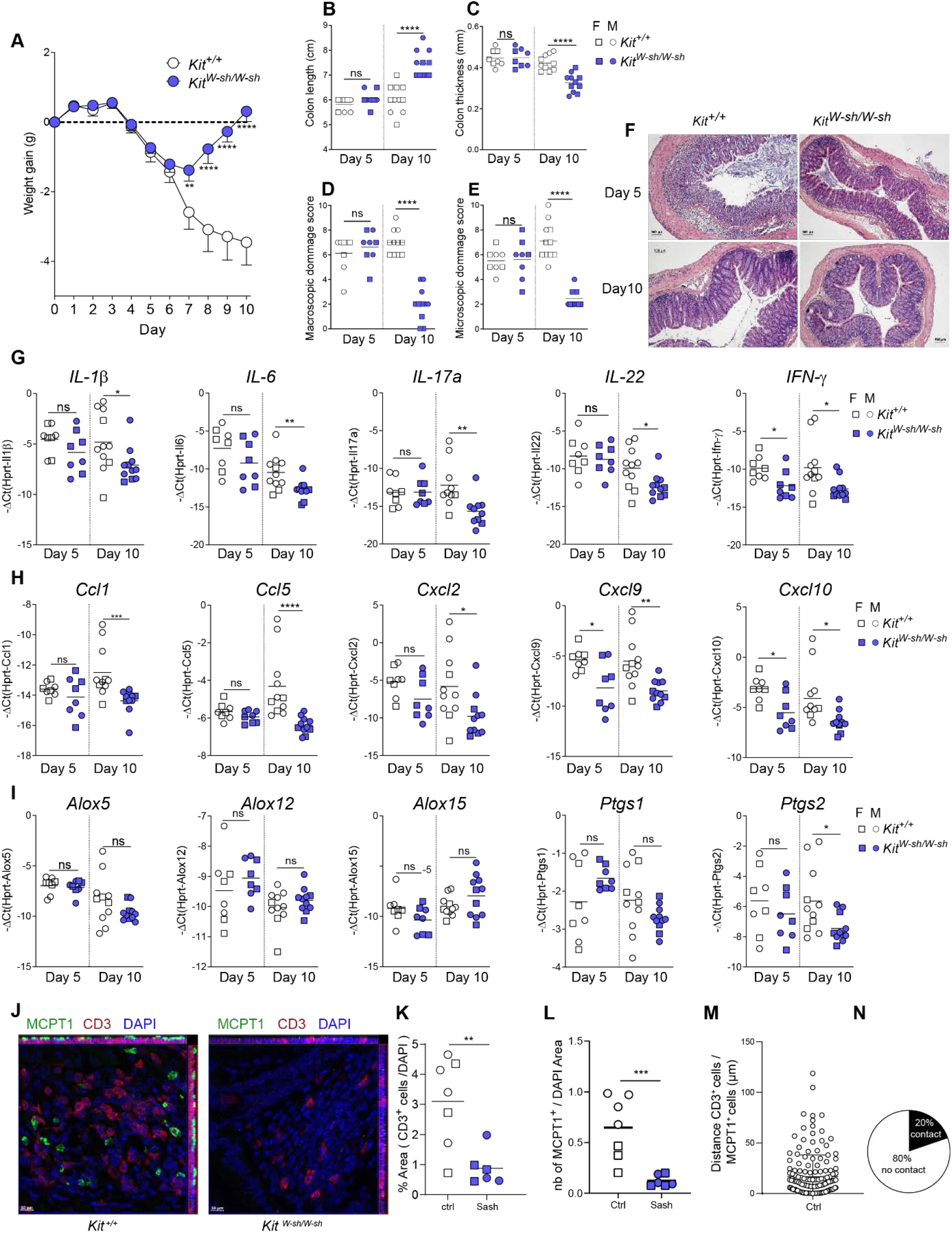
Late-stage colitis severity is reduced in KIT-deficient mice. Colitis was induced by adding 3% DSS into the drinking water for 5 days of MC deficient male and female mice (Kit^W-sh/W-sh^) and their littermate counterpart (Kit^+/+^). (A-F) Weight gain was observed in DSS-treated MC deficient male and female mice (Kit^W-sh/W-sh^, blue circle) and their littermates (Kit^+/+^, white circle). Data are expressed as mean ± SEM. Statistical analysis was performed using repeated-measures 2-way ANOVA and a Bonferroni’s multiple comparisons post hoc tests. *** p<0.001 significantly different from Kit^+/+^ group (A). Colitis severity was assessed by colon length (B), colon thickness (C), macroscopic damage scores (D), microscopic damage scores (E-F). (G-I) Cytokine (G), chemokine (H) and polyunsaturated fatty acid oxygenase (I) gene expression, in DSS-treated MC deficient male (circle) and female (square) mice (Kit^W-sh/W-sh^, blue symbol) and their littermate counterpart (Kit^+/+^, white symbol). Data are expressed as mean ± SEM. Statistical analysis was performed at day 5 or day 10 using Mann-Whitney test. ** p<0.01, *** p<0.001 significantly different from Kit^+/+^ group. (J-N) Colonic tissue samples, recovered at 10 days, of MC deficient mice (Kit^W-sh/W-sh^, right panel) and their littermates (Kit^+/+^, left panel) were stained with anti-CD3 (lymphocytes, red), anti-MCPT1 (mastocytes, green). Cell nuclei were stained with DAPI (blue). Fluorescence images were acquired by confocal microscopy. Scale bar = 10 μm. (J) % area (LTCD3^+^/DAPI) (K) and number of mast cells per area of DAPI determined in the colon of DSS-treated MC deficient male (circle) and female (square) mice (Kit^W-sh/W-sh^, blue symbol) and their littermate counterpart (Kit^+/+^, white symbol) (L). Measure of the distances between randomly selected MCs and their proximal CD3^+^ T cell in the colon of DSS-treated WT (Kit^+/+^) mice (n=157 from 7 mice) (M). Percentage of MCs in direct contact with CD3^+^ T cells in the colon of DSS-treated WT (Kit^+/+^) mice (n=157 from 7 mice) (N).

## Discussion

Providing help to other immune cells was one of the earliest discovered functions of CD4^+^ T cells thereby defining them as “T helper Cell”. First discovered for B cells, it was extended to CD8^+^ T cells and next defined for APCs. Nevertheless, a global analysis, based on deep sequencing, of the help given to other cells was reported only recently for CD8^+^ T cells (Ahrends, 2017 #1638). In this context, it is crucial to investigate the impact of CD4^+^ T cells on tissue-resident immune cells such as MCs, most likely to interact with each other during the development of immune responses. MC are tissular cell uneasy to study due to the difficulty to isolate enough amount of pure cells from human tissues and to the altered functionalities of MCs due to isolation procedures (Lorentz et al., 2015). We employed here peripheral-blood-derived primary human MCs which are widely used and found transcriptionally close to human tissue-resident MCs (Cildir et al., 2019). Here, we revealed that Th cell help alternatively activates MCs by the setting up of a specific transcriptomic program and functional modules such as antigen presentation, inflammatory chemokine and eicosanoid production. In turn, this acquired capacity to dialog with Th cells, allows MCs to mold Th cell responses toward IL-17 production and to induce the emergence of inflammatory Th1.17 cells by a mechanism involving PGE_2_ and IL-1β.

Hypothesizing that CD4^+^ T cells have a major impact on MC responses, here we report that MCs activated by Th cells (helped MCs referred to herein as “MC^TH^”), upregulated a panel of genes allowing them to present antigen via MHC-II molecules (HLA class II molecules expression and enzymatic tools for Ag processing such as Cathepsin S, CD74, HLA-DO, HLA-DM…). More surprisingly, MCs also upregulated several genes involved in Ag cross-presentation and MHC-I molecules. MC cognate interaction with CD8^+^ T cells was previously reported (Stelekati et al., 2009) but remains poorly documented. This phenomenon echoes the help given by Th cell to DCs licensing them with the ability to cross-present Ag to CD8^+^ T cells (Cella et al., 1996; Smith et al., 2004). MC^TH^ also display an inflammatory phenotype defined by their ability to produce CXCR3 family ligands CXCL9 and CXCL10 involved in the recruitment of CXCR3^+^ T cells into the inflammatory site (Karin, 2020), to highly express CXCL2, TNFα, oncostatin M or GM-CSF, to secrete PGD_2_ and IL-1β, to upregulate FcγRI and to reduce their degranulation threshold in response to IgE/Ag stimulation. Even if a complex network of signalization is expected to orchestrate MC^TH^ polarization toward an inflammatory phenotype, the upregulation of genes associated to the response to IFN-γ and the need of close proximity with T cells, argue for a role of IFN-γ and direct contacts. Moreover, in silico analysis suggests that MC^TH^ transcriptional response is triggered by typical master regulatory transcription factors such as IRF8, IRF1, STAT1 and NFKB1 (Platanitis and Decker, 2018). This phenomenon that sounds like macrophage polarization points out MC plasticity.

We then analyzed the impact of MC^TH^ on Teff cell response because T effector/memory cells generated in secondary lymphoid organ (SLO) are recruited in sites of inflammation or recirculate in peripheral organ such as skin or gut lamina propria (Marelli-Berg et al., 2008). Once infiltrated, Teff cell function are induced by either Ag-dependent or Ag-independent stimuli. MC are strategically located near blood vessel allowing them to frequently encounter infiltrating T cells. Thus, tissue-infiltrating Teff cells recently activated via their TCR in SLO may recognize cues from neighboring MC or be reactivated via their TCR by local professional APCs at the vicinity of MCs (Honda et al., 2014b; Wakim et al., 2008) and these later receive help and MC-Th cell interplay can proceed. This phenomenon allows to fine tune T cell activation by distinct innate sentinel cells each providing a specific set of soluble or membrane ligands for T cells. Indeed, we show here that helped MC display a rich panel of costimulatory molecules including ICOSL, CD80, PD-L1, PD-L2, galectin9 and several members of the TNF superfamily molecules.

MCs are often associated to type 2 immune responses but here we show that helped MCs may also be prone to drive Th17 responses and to dampen Treg cells by producing PGE_2_ and IL-1β. Mechanistically, MC^TH^ induced IL-17 production from committed CD4^+^ CCR6^+^ Th cell toward the Th17 profile by producing PGE_2_ and IL-1β. Our results are in line with previous reported showing a pro-Th17 effect of PGE_2_ on Ag-experienced T cells (Boniface et al., 2009; Chizzolini et al., 2008; Napolitani et al., 2009; Yao et al., 2009). IL-1β is a well-known Th17 polarizing factor and its production by MC^TH^ was not dependent of eicosanoid production. IL-1β plays a critical role during the reactivation of previously primed effector and memory CD4+ T cells by triggering cytokine production (Jain et al., 2018). It has recently been reported that conversely T cells also instruct macrophages to produce inflammasome-independent IL-1β (Jain et al., 2020). Here, we extend these results by showing that MCs are also instructed by Teff cells to produce IL-1β. MC^TH^ indeed upregulated genes coding for Caspases 1, 4, 7 and 10, Gasdermin D, FAS, NOD2 and expressed *NLRP3* mRNA (Table S2), suggesting an inflammasome-dependent IL-1β production.

More strikingly we show that MC^TH^ lead to the emergence, via IL1β and PGE_2_, of a Th1.17 cell subset, producing IFN-γ and GM-CSF in combination with IL-17, described as pathogenic in some IMIDs (Hu et al., 2017). Whilst several studies report that IL-23 promotes the differentiation of Th1.17 cells, we show here that these cells can also be induced by MCs independently of IL-23 pointing out the complexity of Th17 regulation. In searching the origin of Th1.17 cells, we identified a Th17 oriented CCR6^+^CCR4^-^CCR10^-^CXCR3^-^ cell population that can be elicited by MCs to produce both IL-17 and IFN-γ in an eicosanoid dependent manner. This population was described as an early stage of the Th17 differentiation maintaining stem cell capability (Muranski et al., 2011; Wacleche et al., 2016).

The capacity of MC^TH^ to promote Th1.17 cells while inhibiting Treg differentiation outlines a detrimental MC-CD4^+^ T cell axis in IBD and probably in still unknown other IMID. Indeed, MCs have been proposed to be involved in IMIDs such as multiple sclerosis, rheumatoid arthritis, type I diabetes or psoriasis (Christy and Brown, 2007; Walker et al., 2012). Ulcerative colitis-associated MCs are transcriptionally affected by T cell-derived IFN-γ and TNF in inflamed tissues (Chen et al., 2021) and *in silico* analysis reveals that a number of genes associated with IBD and other IMIDs overlapped with those upregulated genes in MC^TH^. In Crohn’s disease, MCs and CD4^+^ T cells are in close proximity of each other, highlighting the potential of MCs to promote colitogenic Th17 cells. In addition, our results indicate that MC-CD4^+^ T cell crosstalk plays a key role in development of DSS-induced colitis as it has previously been suspected in experimental allergic encephalomyelitis (Russi et al., 2018; Russi et al., 2016; Secor et al., 2000).

## Material and Methods

### RESOURCES TABLE

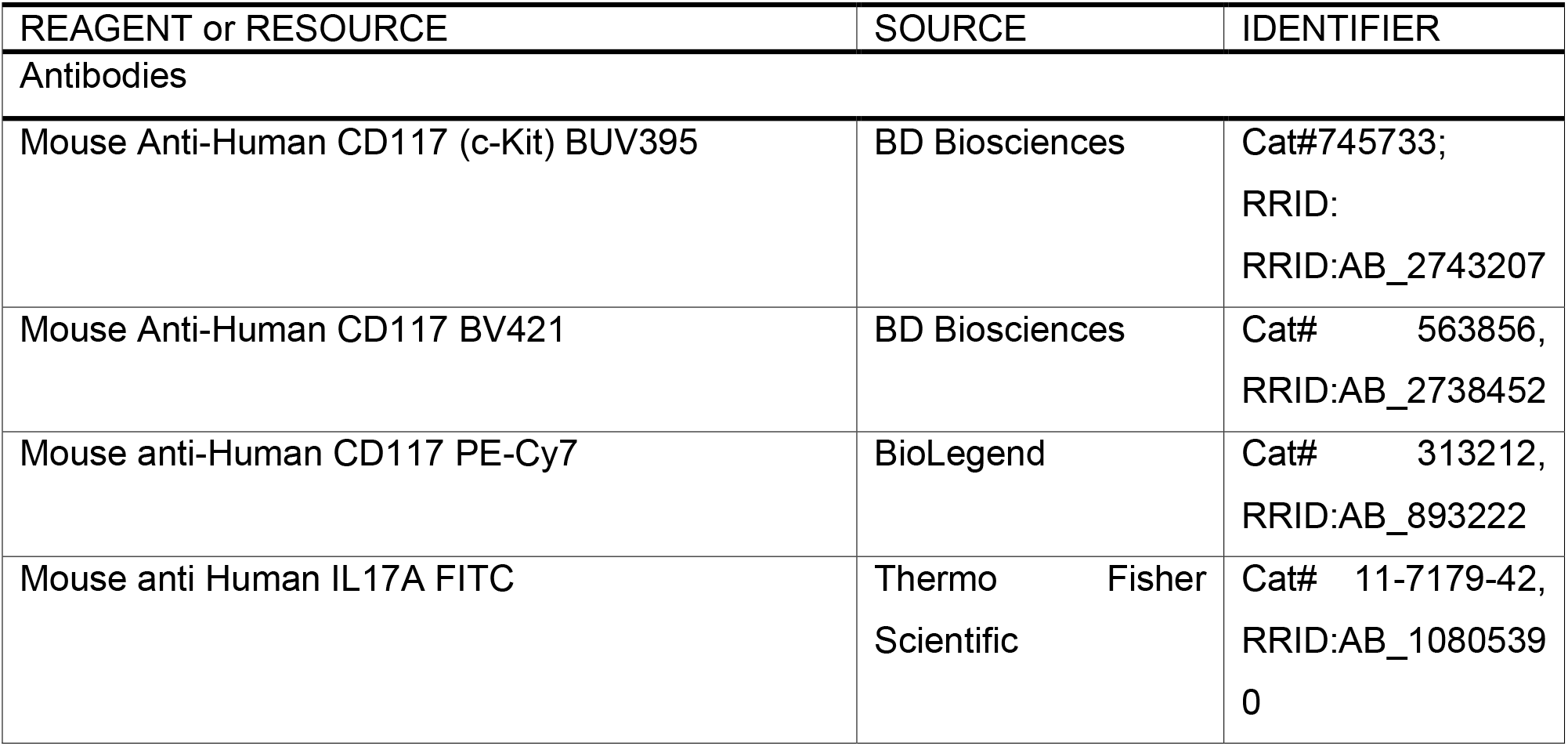
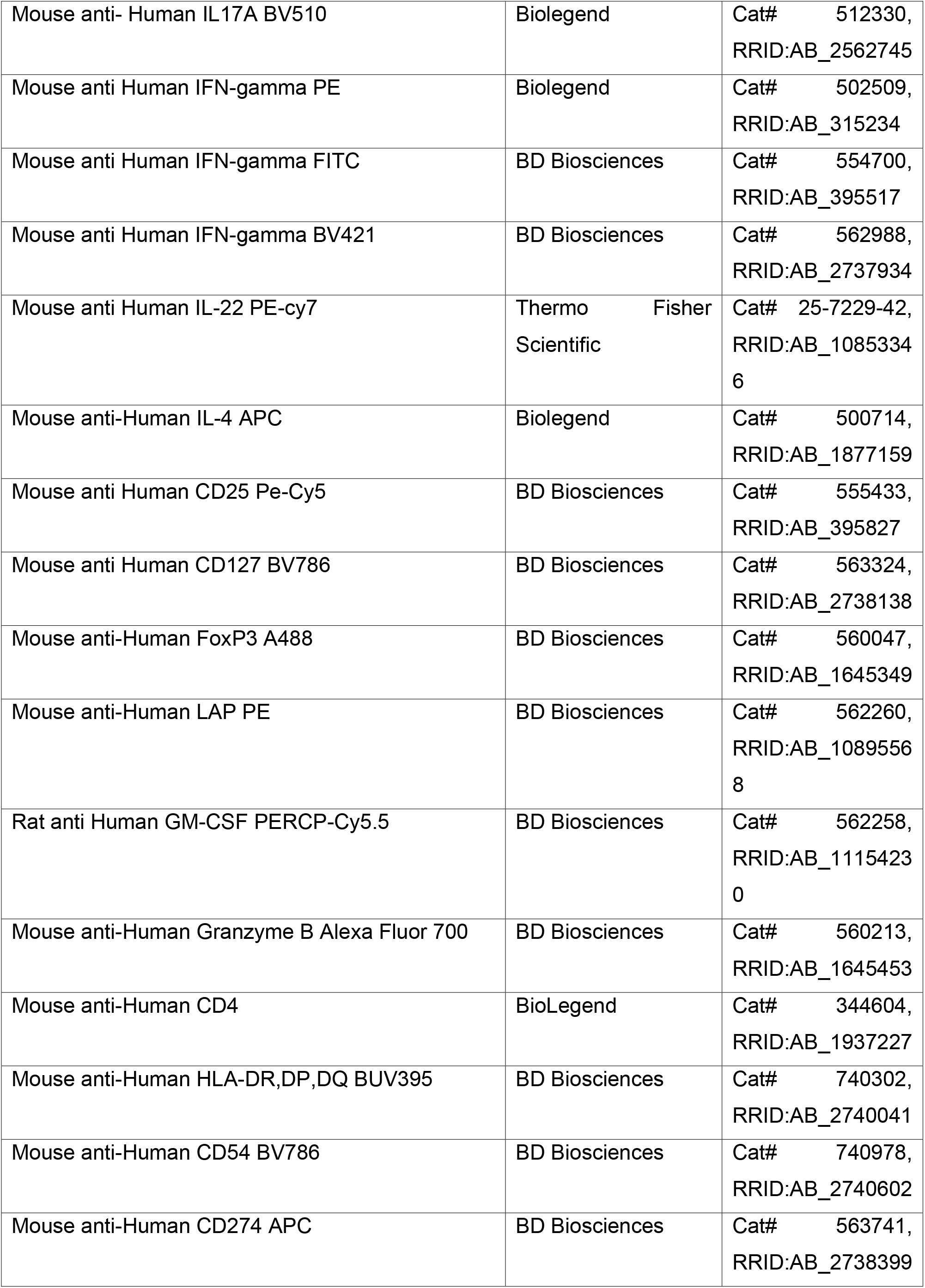
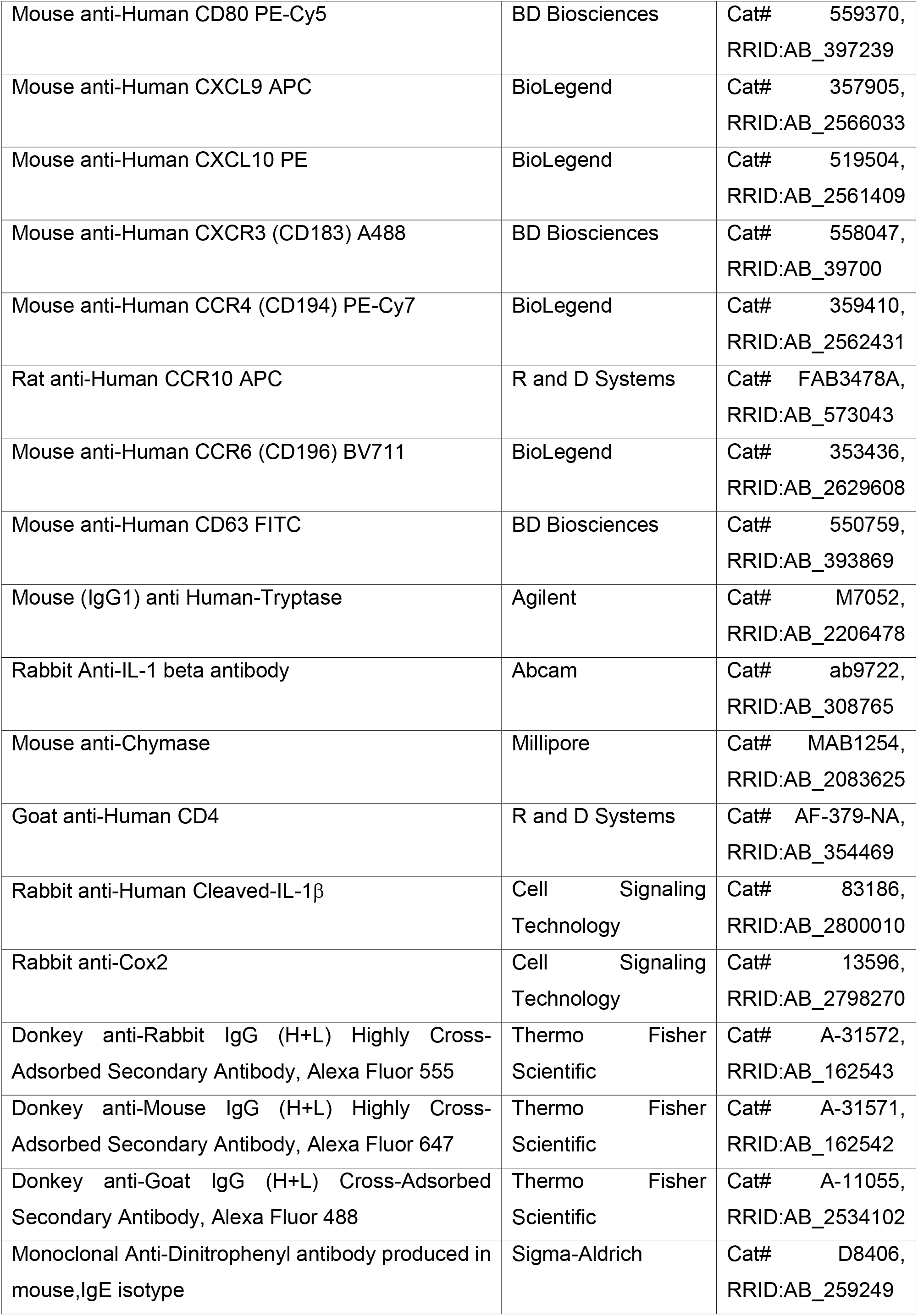
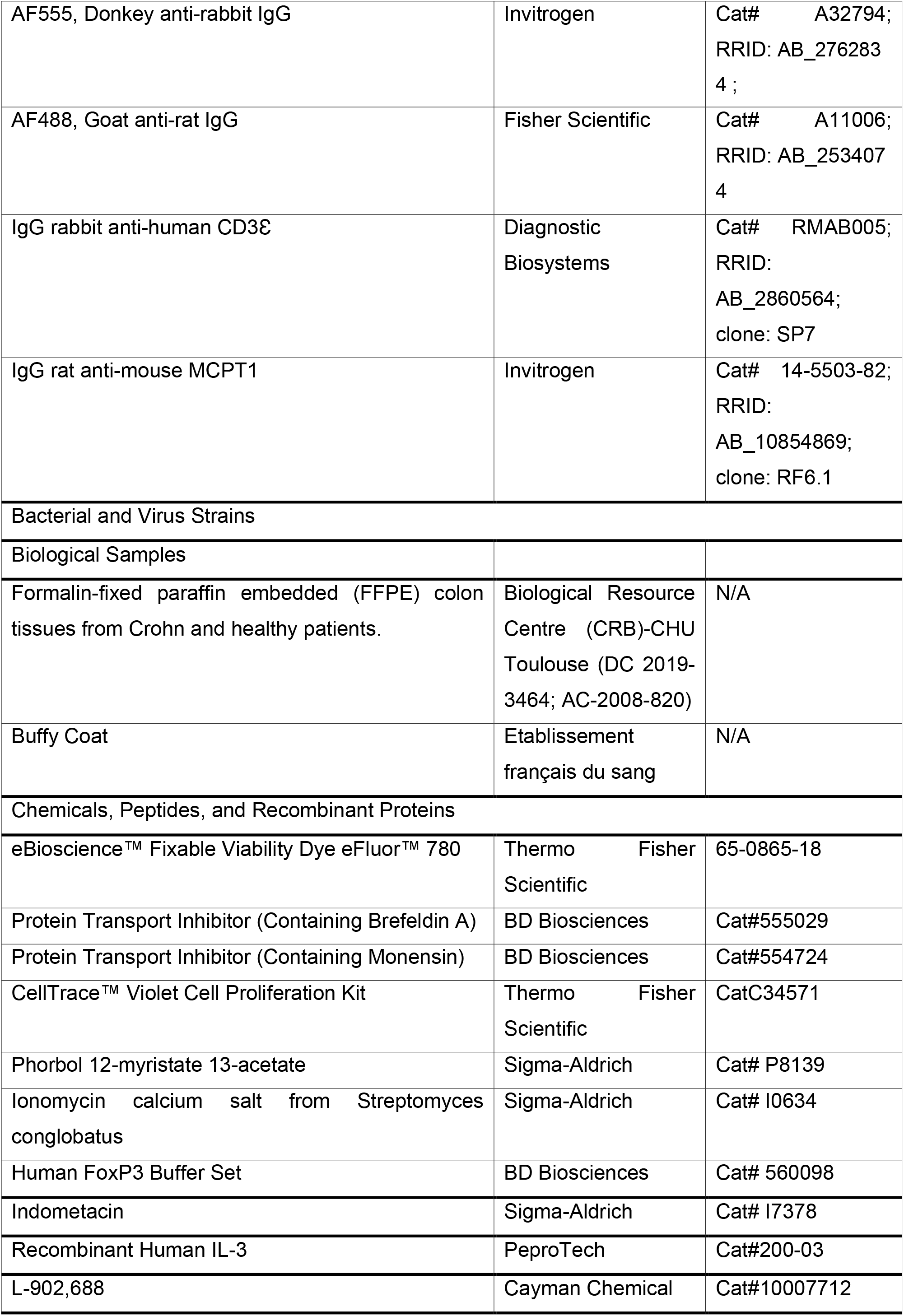
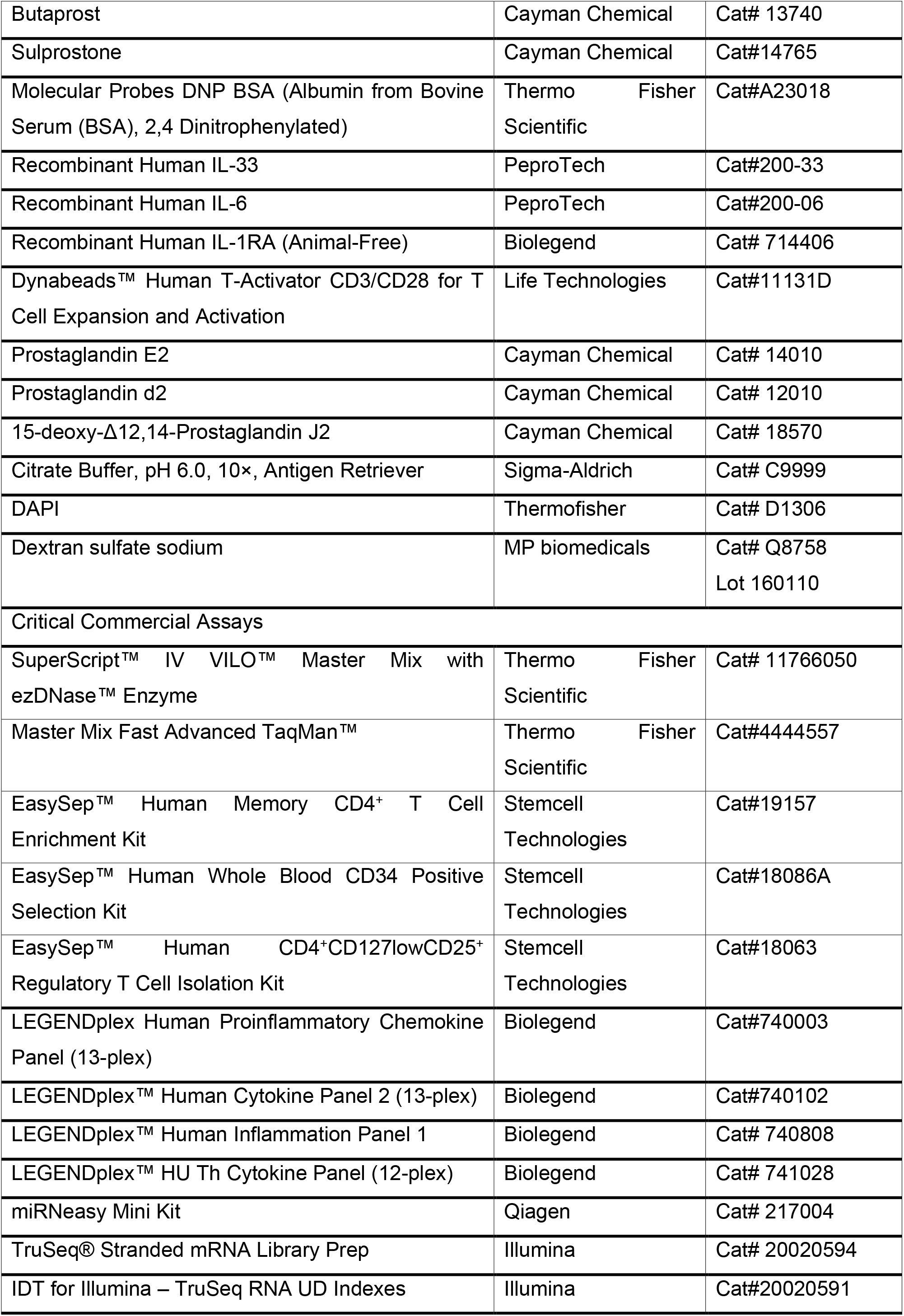
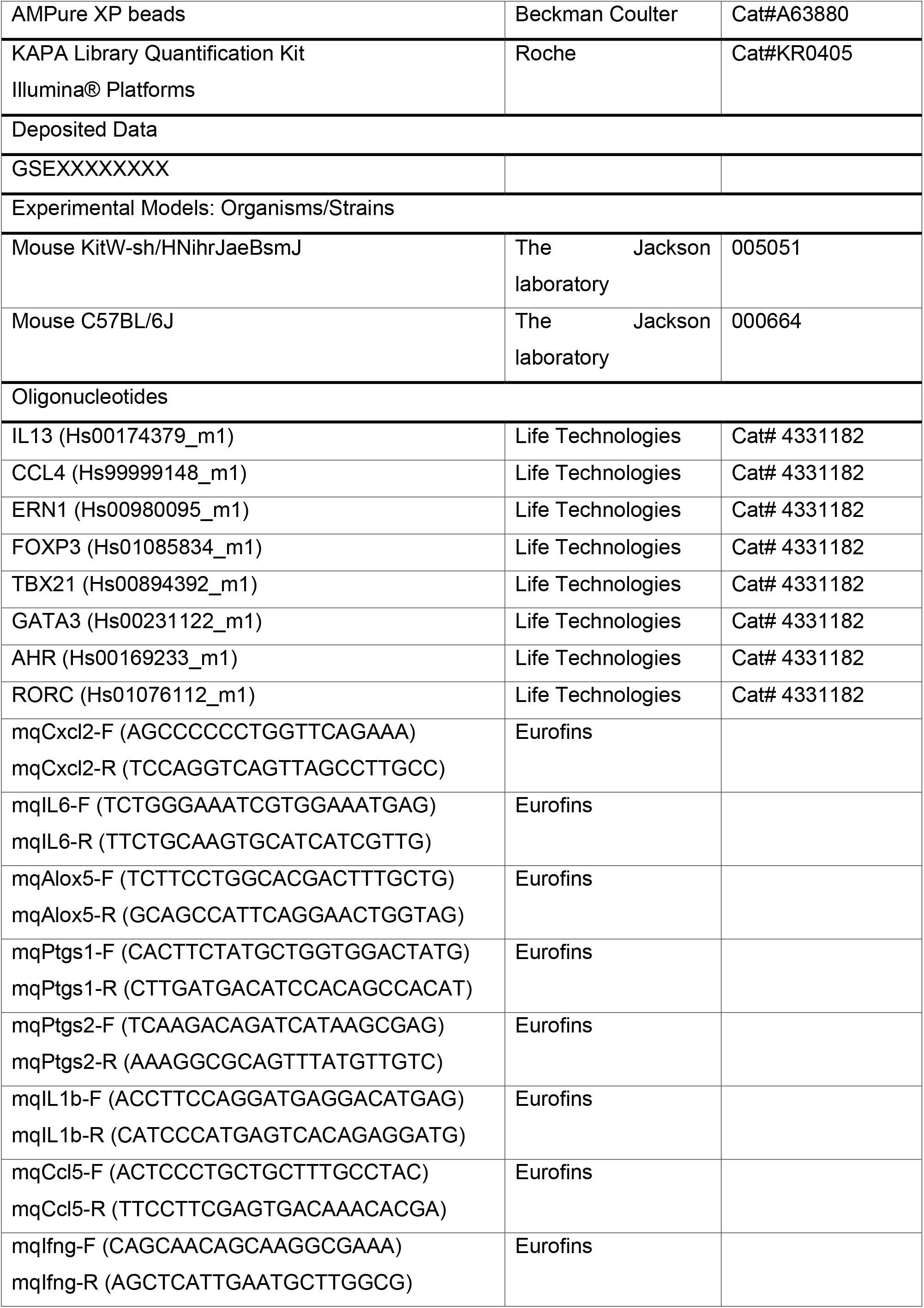
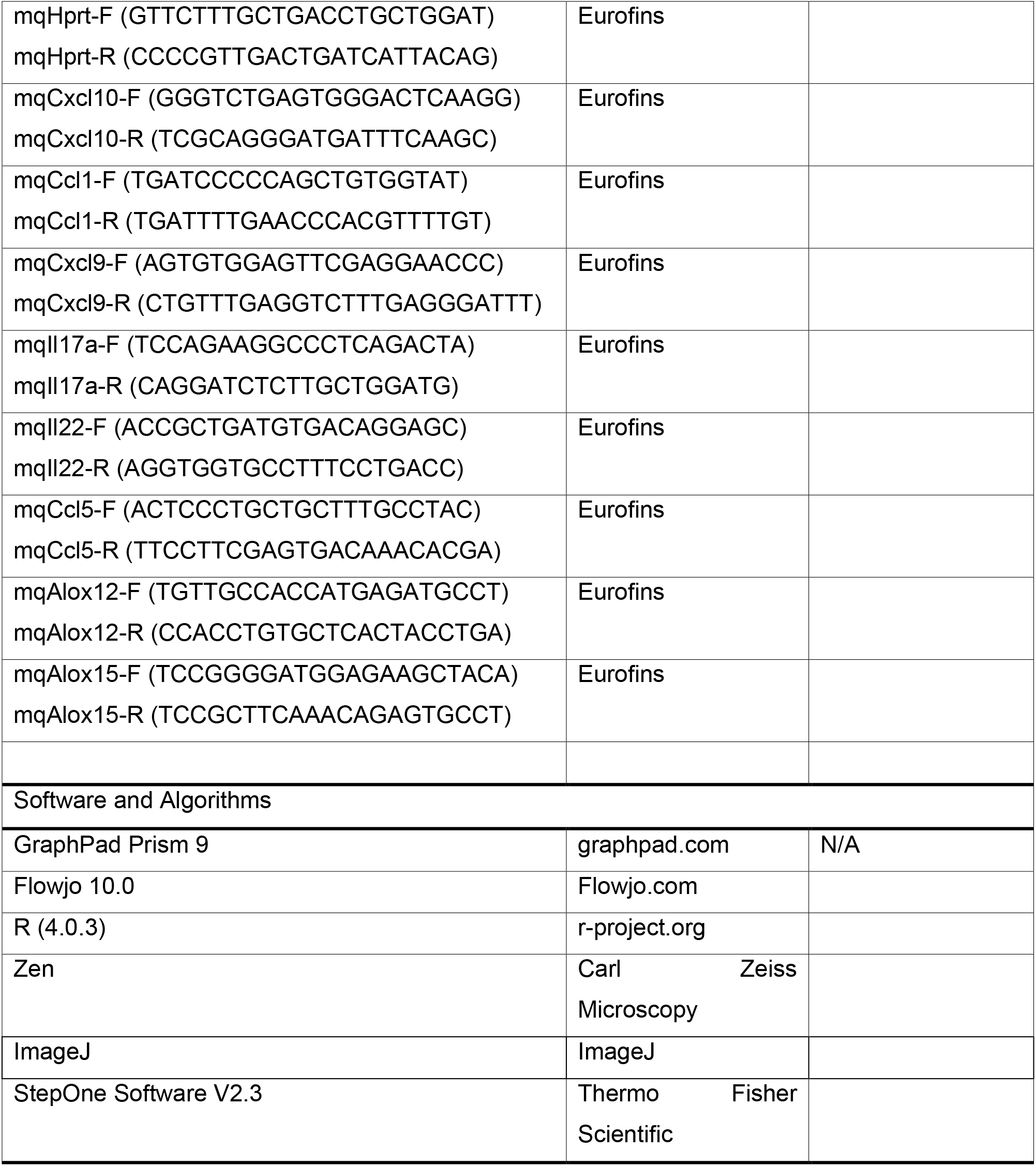

### RESOURCE AVAILABILITY

#### Lead contact

Further information and requests for resources and reagents should be directed to and will be fulfilled by the lead contact, Eric Espinosa.

#### Material availability

This study did not generate new unique reagents.

#### Data and code availability

Processed gene expression data from RNA-sequencing comparisons (Figure 1) are available in supplementary tables, and raw unprocessed files are freely available at GSEXXXXXXX).

### EXPERIMENTAL MODEL AND SUBJECT DETAILS

#### Human primary MC cultures

Peripheral blood mononuclear cells (PBMCs) were obtained from buffy coats (Etablissement Français du Sang). CD34^+^ precursors cells were isolated from PBMCs (EasySep™ Human CD34 Positive Selection Kit, STEMCELL Technologies). CD34^+^ cells were grown under serum-free conditions using StemSpan^TM^ medium (STEMCELL Technologies) supplemented with recombinant human IL-6 (50 ng/mL; Peprotech), human IL-3 (10 ng/mL; Peprotech) and 3% supernatant of CHO transfectants secreting murine SCF (a gift from Dr. P. Dubreuil, Marseille, France, 3% correspond to ∼50 ng/mL SCF) for one week. Cells were grown in IMDM Glutamax I, sodium pyruvate, 2-mercaptoethanol, 0.5% BSA, Insulin-transferrin selenium, Penicillin/Streptomycin (100 U/mL / 100 µg/mL) (all from Invitrogen), IL-6 (50 ng/mL) and 3% supernatant of CHO transfectants secreting murine SCF for 8 weeks and tested phenotypically (Tryptase^+^, CD117^+^, FcεRI^+^) and functionally (β-hexosaminidase release in response to FcεRI crosslinking) before use for experiments. Only primary cell lines showing more than 95% CD117^+^/FcεRI^+^ cells were used for experiments.

#### Human studies

Formalin-fixed paraffin-embedded (FFPE) colon sections from patients with Crohn disease were selected after pathological examination. Normal colon biopsies from patients were used as control. All samples were processed in Biological Resource Center (CRB) of Toulouse, following ethical procedures (Declaration of Helsinki) and written informed consent was obtained from all patients. CRB collections were declared to the Ministry of Research and a transfer agreement was obtained after approval from the appropriate ethics committees (DC 2019-3464; AC-2008-820).

Multi-omics data were collected from the Open Targets Platform (https://www.targetvalidation.org/).

#### Animals

C57BL/6J (Kit^+/+^) and KitW-sh/HNihrJaeBsmJ (Kit^W-sh/W-sh^) mice aged 10-14 weeks (The Jackson laboratories) were used. Mice were raised in sanitary conditions without pathogens with free access to water and food and submitted to alternating cycles of 12 hours of light and darkness. All procedures were performed in accordance with the Guide for the Care and Use of Laboratory Animals of the European Council and were approved by the Animal Care and Ethics Committee of US006/CREFE (CEEA-122; application number APAFIS #7762-CE2016112509278235V2).

### METHOD DETAILS

#### coculture experiments

1 × 10^5^ memory CD4^+^ T cells freshly purified from PBMC by negative selection and magnetic separetion(EasySep™ Human memory CD4^+^ T cell enrichment kit, STEMCELL Technologies) were cocultured at 1:1 ratio for 6 days with anti-CD3/CD28–coated beads (Dynabeads, Life Technologies) at 1 bead: 10 T cells in RPMI 1640 supplemented with 10% serum replacement medium (knockout medium, Life Technologies), GlutaMAX-I, sodium pyruvate, 2-mercaptoethanol, and 1% supernatant of CHO transfectants secreting murine SCF. Coculture were treated as indicated with either indomethacin (100 µmol/L, Sigma-Aldrich), PGE_2_ (11-300 ng/mL, Cayman Chemical), PGD2 (11-300 ng/mL, Cayman Chemical), 15d-PGJ2 (11-300 ng/mL, Cayman Chemical), Butaprost (10 µmol/L, Cayman Chemical), Sulprostone (10 nmol/L, Cayman Chemical), L902,688 (12 nmol/L, Cayman Chemical), IL-1Ra (100 ng/mL, Biolegend). For proliferation assay memory CD4^+^ T cells were stained with CellTrace™ Violet Cell Proliferation Kit (Thermo Fisher Scientific) according to the Manufacturer’s recommendations. When employed, Treg were purified from CD4^+^ memory T cells by magnetic separation (EasySep™ Human CD4^+^CD127^low^CD25^+^ Regulatory T Cell Isolation Kit, STEMCELL Technologies) and labelled with CellTrace™ Violet. CTV-Labelled Treg cells were reincorporated with their non-Treg counterparts.

#### MC degranulation assay

MCs were sensitized with anti-DNP human IgE (1 µg/mL, Sigma-Aldrich) for 16 hours and next cocultivated with memory CD4^+^ T cells (ratio 1:1) with or without anti-CD3/28 coated beads for 48 hours. Cells were washed in Tyrode’ Buffer and distributed in 96-well flat-bottom plate in 50 µL Tyrode’s buffer and adapted to 37°C for 20 minutes. hMCs were then treated with DNP BSA (0.16-20 ng/mL, Thermo Fisher Scientific) for 45 min at 37°C, 5% CO_2_. Cells were next stained with anti-CD63-FITC mAb (H5C6) at 4°C for 30 min and proceeded to flow cytometry analysis.

#### PUFA metabolites LC-MS/MS measurements

PUFA metabolites were extracted from the supernatant or the cell pellets following the methods previously described (Le Faouder et al., 2013). 6-keto-prostaglandin F1 alpha (6kPGF1α), thromboxane B2 (TxB2), PGE_2_, 8-isoPGA_2_, PGE_3_, 15d-PGJ_2_, PGD_2_, lipoxin A4 (LxA4), LxB4, 7-Maresin 1 (7-Mar1), leukotriene B4 (LtB_4_), LtB_5_, protectin Dx (PDx), 18-hydroxyeicosapentaenoic (18-HEPE), 9-hydroxyoctadecadienoic acid (9-HODE), 13-HODE, 15-hydroxyeicosatetraenoic acid (15-HETE), 12-HETE, 8-HETE, 5-HETE, 17-hydroxydocosahexaenoic acid (17-HDoHE), 14-HDoHE, 14,15-epoxyeicosatrienoic acid (14,15-EET), 11,12-EET, 8,9-EET, 5,6-EET, and 5-oxoeicosatetraenoic acid (5-oxoETE) were quantified. To simultaneously separate 27 lipids of interest and three deuterated internal standards (5-HETEd8, LxA4d4 and LtB4d4), LC-MS/MS analysis was performed on an ultrahigh-performance liquid chromatography system (UHPLC; Agilent LC1290 Infinity) coupled to an Agilent 6460 triple quadrupole MS (Agilent Technologies) equipped with electrospray ionization operating in negative mode. Reverse-phase UHPLC was performed using a ZorBAX SB-C18 column (Agilent Technologies) with a gradient elution. The mobile phases consisted of water, acetonitrile (ACN), and formic acid (FA) [75:25:0.1 (v/v/v)] (solution A) and ACN and FA [100:0.1 (v/v)] (solution B). The linear gradient was as follows: 0% solution B at 0 min, 85% solution B at 8.5 min, 100% solution B at 9.5 min, 100% solution B at 10.5 min, and 0% solution B at 12 min. The flow rate was 0.35 ml/min. The autosampler was set at 5°C, and the injection volume was 5 µL. Data were acquired in multiple reaction monitoring (MRM) mode with optimized conditions. Peak detection, integration, and quantitative analysis were performed with MassHunter Quantitative analysis software (Agilent Technologies). For each standard, calibration curves were built using 10 solutions at concentrations ranging from 0.95 to 500 ng/ml. A linear regression with a weight factor of 1/X was applied for each compound. The limit of detection (LOD) and the limit of quantification (LOQ) were determined for the 28 compounds using signal-to-noise (S/N) ratios. The LOD corresponded to the lowest concentration leading to an S/N value >3, and LOQ corresponded to the lowest concentration leading to an S/N value >10. All values less than the LOQ were not considered. Blank samples were evaluated, and their injection showed no interference (no peak detected), during the analysis.

#### Flow cytometry and cell sorting

Cell surface staining with fluorochrome-labelled primary Abs were performed in PBS 1% FCS 1% human serum (HS) at the concentration recommended by the manufacturer at 4°C for 30 min. Cell viability was ascertained by labeling with fixable viability dyes (eBioscience).

For T cell cytokine intracellular staining, cells were stimulated with PMA (50 ng/mL) and Ionomycin (1 µg/mL) in presence of GolgiPlug™ and GolgiStop™ (BD biosciences) for 5 hours. After washing in PBS, cells were incubated with anti-CD117 (104D2) and viability dye (eBioscience™ Fixable Viability Dye eFluor™ 780) in PBS 1% FCS 1% HS at 4°C for 30 min. Cells were washed in PBS and next fixed in PFA 4% for 10 min. at RT and permeabilized with PBS 1% FCS 1% HS 0.1 % saponin (permeabilization buffer) for 10 min. Cells were next incubated with indicated Abs : anti-IL-17A (BL168), anti-IFN-γ (B27), anti-IL-4 (8D4-8), anti-IL-22 (22URTI), anti-granzyme B (GB11), anti-GM-CSF (BVD2-21C11) in permeabilization buffer for 45min at RT. Cells were next washed in PBS and proceeded to flow cytometry analysis.

Treg cells in cocultures were identified by cell surface staining with anti-CD117 (104D2), anti-CD127 (HIL-7R-M21), anti-CD25 (M-A251), anti-LAP (TW4-2F8) and viability dye (eBioscience™ Fixable Viability Dye eFluor™ 780) followed by Foxp3 staining with anti-FoxP3 (259D/C7) using Human FoxP3 buffer set (BD biosciences) according to the Manufacturer’s recommendations.

For intracellular CXCL9 and CXCL10 staining, MC and CD4^+^ effector/memory T cells were cocultured for 48 h and GolgiPlug™ was added 4 hours before the end of the coculture. Cells were washed in PBS and proceeded for cell surface staining with anti-CD117 (104D2) and viability dye (eBioscience™ Fixable Viability Dye eFluor™ 780) in PBS 1% FCS 1% HS at 4°C for 30 min. Cells were washed and Intracellular staining with anti-CXCL9 (J1015E10) and CXCL10 (J034D6) was carried out as indicated above.

For T helper cell subset sorting, effector/memory CD4^+^ T cells were labeled with anti-CXCR3 (1C6/CXCR3), anti-CCR4 (L291H4), anti-CCR6 (G034E3), anti-CCR10 (#314305) and viability dye (eBioscience™ Fixable Viability Dye eFluor™ 780). Cell sorting was performed with indicated gating strategy by using a custom configuration FACSAria cell sorter.

All flow cytometry experiments were acquired using a custom configuration Fortessa flow cytometer and the FACS Diva software (BD Biosciences) and analyzed with FlowJo V10.4.2 software (TreeStar). Cell sorting was performed by using a custom configuration FACSAria cell sorter (BD Biosciences) or FACSMelody cell sorter (BD biosciences).

#### Cytokine quantitation

For CXCL-9 / CXCL-10 / IL-6 / TNF-α / IL-1β / IL-23 / IL-12 / IL-1α Supernatant of coculture were collected after 48h and store at −80°C. For evaluated T cell production of IFN-γ / IL-4 / IL-17 /IL-22, CD4^+^ T cells were sort by FACS at day 6 and restimulate with PMA (50ng/mL) and Ionomycin (1 µg/mL) during 5 hours and supernatant were recovered and store at −80°C. Supernatant concentration were measured by a bead-based multiplex assay (LEGENDplex™, Biolegend) following the Manufacturer’s recommendations. Flow cytometric analysis were performed with a MACSQuant^®^ Analyzer 10 instrument (Miltenyi Biotec).

#### RNAseq Libraries Preparation

750 000 MCs were either FACS-sorted after 24h of coculture with resting or anti-CD3/28 coated beads-activated effector/memory CD4^+^ T cells or stimulated for 4 hours with 10 ng/mL of IL-33 (Peprotech) or with 100 ng/mL of DNP-BSA (thermo-fischer scientific). RNA was extracted using miRNeasy Mini Kit (Qiagen). Libraries were prepared using 1 µg of high-quality total RNA. TruSeq stranded mRNA Library prep kit (Illumina) and Truseq RNA Unique Dual Indexed Adapters (Illumina) are used according to the manufacturer protocol. Size selection was performed using AMPure XP beads (Beckman Coulter). Library size and quality were confirmed on Fragment analyzer (Agilent). Libraries were quantified by qPCR using the KAPA quantification kit for Illumina platforms (KAPA Biosystems, Roche). Samples were pooled in equimolar fashion. The libraries were sequenced on a NextSeq 550 (Illumina) in pair end sequencing 2 × 75 bp with dual indexes (i7 and i5, 8bp).

#### RNAseq Analysis

Read alignment and gene quantification for RNA-seq: FastQ files were generated via llumina bcl2fastq2 (version 2.20.0.422), aligned and quantified using RSEM (RSEM 1.3.0, bowtie2-2.3.4) and the homo sapiens transcriptome reference GRCh38 (v97) (Li and Dewey, 2011). Differential analyses of RNA-Seq data at the gene level were performed using the DESeq2 R package (v1.30.0) with the recommended workflow (Love et al., 2014). ranked DEG lists (FDR<0.01 and fold-change>1.5) were established using the *apeglm* method for effect size shrinkage (Zhu et al., 2018). Venn diagrams were generated using eulerr R package. DEG adjusted p-values (FDR) and log_2_ fold changes were displayed as volcano plots using EnhancedVolcano (v 1.8.0) R package. For visualization and clustering (PCA, heatmaps of sample-to-sample distances), regularized logarithm (rlog) transformation of the count data was performed to remove the dependence of the variance on the mean. R Pheatmap package (v1.0.12) was employed to display DEGs (hierarchical clustering using the euclidean distance and complete linkage method). Heatmaps for specific groups of genes were generated using R package ComplexHeatmap (v 2.6.2) to visualize both genes clustered by expression and log_2_foldchange plus TPM values (Gu et al., 2016). We performed Reactome pathway enrichment with the Bioconductor package ReactomePA (Yu and He, 2016) of DEGs between MC +T_ACT_ and MC +T_CTRL_ conditions (FDR<0.05).

#### Transcription factor target enrichment

Transcription factor target enrichment among upregulated genes (FDR < 0.01 and fold change>2) in stimulated vs unstimulated MCs was performed with Cytoscape (v 3.8.2) in combination with the iRegulon plugin(Janky et al., 2014) using the Homo sapiens database and default settings. Transcription factors of interest with motif enrichment scores (NES) > 3 were kept.

#### Intercellular communication score computation

To infer helped MC-activated CD4^+^ memory T cell communication network, we based our analysis on publicly available database of ligand-receptors pairs collated by Noel and co-workers (Noel et al., 2021) and on computation method previously described (Armingol et al., 2021; Noel et al., 2021). Thresholding method was used to infer “active” L-R pairs and communication score computation (expression product method) was used to rank the active L-R pairs. Briefly, Average expression levels (FPKM) of the genes corresponding to the ligands in the database were selected from the list of DEGs between helped and non-helped mast cells (FDR<0.01 and log_2_ fold change >1) (cf RNAseq analysis above). Average expression levels (mean FPKM value) of the genes corresponding to the receptors in the database were selected from RNAseq data of memory CD4+ T cells activated with anti-CD3/CD28 coated beads for 24h (GEO accession number GSE73214) (LaMere et al., 2017). For Thresholding, only ligands and receptors with FPKM>1 were kept in the analysis. Intercellular communication score was calculated as previously described (Noel et al., 2021): ligand and receptor gene expression were next scaled by the maximum value of gene expression among ligands and receptors respectively and multiplied by 10 to get values ranging from 0 to 10. The maximum value of gene expression was calculated as the mean of the 10% highest values. Scaled values above 10 (outliers) were coerced to 10. Because the database employed here takes into account multichain receptors or ligands, we first calculated the expression level for the multichain molecules as the geometric average of the values of the ligand /receptor chain expression. Next the ligand-receptor pair score was determined by the product of the expression level of the ligand by the expression level of the receptor (if L_i_ is the average expression level of ligand i by helped MC and R_i_ is the is the average expression level of the corresponding receptor by activated CD4^+^ memory T cell, the score S_i_ of this interaction is S_i_=L_i_.R_i._ Inferred active Ligand-receptor pairs (S_i_>0) are provided in Table S2. Custom R scripts were used for analyses. For circular visualization of the links between MC ligands and activated memory CD4^+^ T cell corresponding receptors, the circlize R package was used (Gu et al., 2014) to represent the 30 top-ranked L-R pairs according to their computed communication score.

#### Overlap between MC^TH^ upregulated genes and disease-associated genes

We used the Open Targets Platform (Ochoa et al., 2020) (https://platform.opentargets.org/) to obtain a ranked list of target genes associated with a pathology. This platform integrates validated evidences from genetics, genomics, transcriptomics, drugs, animal models and scientific literature (GWAS, pheWAS catalog, Uniprot…) to score and rank target-disease associations (Ochoa et al., 2020). We chose several IMID and non-IMID (e.g. infectious diseases and cancers) and obtained associated gene list ranked by an association score ranging from 0 to 1. Target genes with association score>0.5 were kept for the overlap analysis).

MC^TH^ upregulated genes and disease-associated gene lists were compared using the GeneOverlap R package (v 1.26.0). GeneOverlap performs Fisher’s exact test to calculate the significance of each pair of gene lists in comparison to a genomic background (21,196 human protein-coding genes). The function returns the number of common genes between the two lists, the P value, and the estimated odds ratio. Odds ratio larger than 1 indicate a positive association between lists. The overlap between lists was represented by plotting odds ratios and P values for each disease. For IBD, we also used a curated prioritized gene list from GWAS previously reported (Mukhopadhyay et al., 2019) to calculate overlap with MC^TH^ using GeneOverlap.

#### Quantitative real-time PCR in MC/T cell cocultures

RNA extracted for RNAseq Libraries preparation was also employed for RT-qPCR analysis (RNAseq validation). For transcription factor analysis, total RNA was extracted from 200 000 T cells (sorted at day 6 of coculture using phenol chloroform protocol and RNA concentration was determined using a Clariostar multi-mode plate reader (BMG Labtech). Reverse transcription of total RNA was performed using the SuperScript™ VILO™ cDNA Synthesis Kit according to the manufacturer’s recommendations (thermo Fisher scientific). real-time PCR was performed with Master Mix Fast Advanced TaqMan™ (Thermofisher Scientific) using a StepOnePlus™ Real-Time PCR detection system (Applied Biosystems™). Following primers were used: IL13 (Hs00174379_m1), CCL4 (Hs99999148_m1), ERN1 (Hs00980095_m1), TBX21 (Hs00894392_m1), GATA3 (Hs00231122_m1), FOXP3 (Hs01085834_m1), AHR (Hs00169233_m1), RORC (Hs01076112_m1). Gene expression was normalized as n-fold difference to the housekeeping gene GAPDH (Hs03929097_g1) based on the 2^-ΔΔCt^ method. Relative quantification of gene expression was performed with the StepOne Software v2.3 (Applied Biosystems™).

#### Immunofluorescence and confocal microscopy

MC and CD4+ memory T cells were cocultured with or without anti-CD3/CD28 coated beads for 48 h. Cells were plated on Poly-D-Lysine (Sigma) coated slides and were fixed in PFA 4% for 10 min at RT and permeabilized with PBS 1% BSA 0.1% saponin for 5 minutes. Cells were stained with mouse anti-chymase (IgG1k, clone B7), goat anti-CD4 (polyclonal Ab) and rabbit anti-IL-1β (polyclonal Ab) at RT for 1h in PBS 1% BSA 0.1% saponin. Cells were next incubated with secondary antibodies: donkey anti-rabbit IgG (H+L) Alexa Fluor 555 (polyclonal), donkey anti-mouse IgG (H+L) Alexa Fluor 647 (polyclonal), donkey anti-goat IgG (H+L), Alexa Fluor 488 (polyclonal). Cells were next counterstained with DAPI (1 µg/mL).

For IF studies on human colon biopsies, 3µm thick sections of FFPE tissues were provided by the Biological Resource Centre of the CHU Toulouse (CRB). Sections were first rehydrated in successive bath of: Xylène (20 min.) / 100% ethanol (20 min.) / 95% ethanol (5min.) / 70% ethanol (5 min.) / 50% ethanol (5min.) / PBS (5 min.). Antigen retrieval protocol were achieved using Citrate buffer pH 6 (Sigma-Aldrich). Sections were saturated during 30 minutes in PBS 10% Human Serum 0,3% Saponin and incubated at 4°C with primary antibody overnight in PBS 10% Human Serum 0,3% Saponin. Sections were next washed 4 times and exposed to secondary antibody for 2 hours at RT. Before covering, samples were washed three times and incubated for 5 min with DAPI (1 µg/mL). The following primary antibodies were used: Mouse (IgG1) anti Human-Tryptase (Clone AA1), Goat anti-Human CD4 (polyclonal), Rabbit anti-Human Cleaved-IL-1β (D3A3Z), Rabbit anti-Cox2 (D5H5). The following secondary antibodies from Thermo Fischer Scientific were used: Donkey anti-Rabbit IgG (H+L) Highly Cross-Adsorbed Secondary Antibody, Alexa Fluor 555 (polyclonal), Donkey anti-Mouse IgG (H+L) Highly Cross-Adsorbed Secondary Antibody, Alexa Fluor 647 (polyclonal), Donkey anti-Goat IgG (H+L) Cross-Adsorbed Secondary Antibody, Alexa Fluor 488(polyclonal). Images were acquired using Zeiss LSM 780 confocal microscope. Images acquired were analyzed using Zen (Carl Zeiss Microscopy) and Image J software.

#### Dextran Sulfate Sodium–Induced Colitis

Colitis was induced by adding 3% (weight/volume) dextran sulfate sodium (DSS) to the drinking water for 5 days. Then, from day 5 to day 10, animals received only water.

#### Macroscopic Assessment of Inflammation-Associated Colon Damage

Macroscopic colonic tissue damage was evaluated using a scale ranging from 0 to 11 as follows: erythema (0, absent; 1, length of the area <1 cm; 2, length of the area >1 cm), edema (0, absent; 1, mild; 2, severe), strictures (0, absent; 1, one; 2, two; 3, more than two), ulceration (0, absent; 1, present), mucus (0, absent; 1, present), and adhesion (0, absent; 1, moderate; 2, severe). Colon wall thickness was measured with an electronic caliper.

#### Histological Assessment of Inflammation-Associated Colon Damage

Colonic tissue specimens were excised 2 cm distal to the anus and immediately transferred into 10% formaldehyde to be embedded in paraffin. Five-micrometer colonic sections were then stained with H&E. Slides were examined and graded for cellular infiltration, mucosal architecture alteration and submucosal edema from 0 to 3 (absent, mild, moderate and severe) and vasculitis, muscular thickening, crypt abscess and goblet cell depletion from 0 to 1 (absent or present); the maximal score being 13.

#### Real time PCR analysis in mouse colon

Colon biopsies were crushed in 500 µL of Trizol (Molecular Research Center, Euromedex, Souffelweyersheim, France) in Precellys lysing kit tubes (Bertin Technologies, Montigny le Bretonneux, France) placed in a Precellys (2823g, 30 seconds twice; Bertin Technologies). After addition of chloroform and centrifugation (15 minutes, 1019 g at 4°C), the supernatant containing the RNA was removed. Ethanol 70% (vol/vol) was added and the contents were placed in columns (GenElute® mammalian total RNA miniprep kit, Sigma Aldrich). The RNA was extracted according to the manufacturer recommendations. Subsequently the RNA were dosed in a nanodrop (implenGmbH, Dominique Dutscher, Issy-les-Moulineaux, France). RNA (10 µg) preparations from colon of DSS mice were purified using the kit (Dynabeads® mRNA purification kit, Fisher Scientific, Illkirch France) according to the manufacturer recommendations. Then, total RNA was reverse-transcribed with Moloney murine leukemia virus reverse transcriptase (Fisher Scientific) using random hexamers (Fisher Scientific) for priming. Transcripts encoding hypoxanthine phosphoribosyl transferase (Hprt), chemokine (C-C motif) ligand 1 (Ccl1), Ccl5, interleukin 6 (Il6), Il1β, Il17A, Il22, chemokine (C-X-C motif) 2 ligand (Cxcl1), Cxcl9, Cxcl10, interferon gamma (Ifng), Arachidonate 5-Lipoxygenase (Alox5), Alox12, Alox15, prostaglandin-Endoperoxide Synthase 1 (Ptgs1) and Ptgs2 were quantified by real-time PCR using specific forward and reverse primers (See key resource table for sequences), the kit Takyon No ROX SYBR 2X MasterMix blue dTTP (Eurogentec) and the LightCycler480II (Roche Diagnostics, Meylan, France).

#### Immunofluorescent staining of lymphocytes and mast cells in mouse colon tissue

Colonic sections (5 µm) were saturated with PBS 1% BSA and then incubated with rabbit anti-CD3 (Clone SP7, Diagnostic BioSystems) monoclonal antibody (mAb) and rat anti-mouse MCPT1 (invitrogen) overnight at 4°C. After washing with PBS, bound antibodies were revealed with Alexa Fluor 555-labeled goat antirabbit IgG antibodies (Invitrogen) and Alexa Fluor 488-labelled Goat anti-rat. Slides were mounted and nuclei were stained with 4’,6-Diamidino-2-Phenylindole (DAPI) fluorescent mounting medium (VECTASHIELD®, Vector laboratories Inc). Images were taken using confocal laser scanning microscope LEICA TCS SP8 (LEICA microsystems) with ×20 objective. Number of mast (MCPT1^+^) has been determined on 3 different areas per slide and expressed has the number of mast cells per DAPI area. For T cells, the surface of CD3-immunoreactivy has been determined using ImageJ software and expressed as the % of area CD3-IR/ area of DAPI labelling.

### QUANTIFICATION AND STATISTICAL ANALYSIS

#### Statistical Analysis

Statistical tests were performed with Graph Pad Prism V9 software (GraphPad Software, Inc.). Test performed are indicated in the figure legends. Non parametric tests were performed when group distribution failed normality or variance homogeneity. All p values are two-sided, (* p < 0.05; ** p < 0.01; *** p < 0.001; **** p < 0.0001 and ns, not significant).

## Acknowledgments

This work was supported by grants from the Laboratoire d’Excellence Toulouse Cancer (TOUCAN) (contract ANR11-LABEX) and from the Ligue Nationale contre le Cancer (Equipe labellisée 2018). E.L. was supported by fellowships from the French Ministry of Education and Research and from Ligue Nationale contre le Cancer. N.C is a recipient of the grant from ANR (ANR-18-CE14-0039-01). We thank the CRCT and Infinity core facilities for assistance. We gratefully acknowledge animal care facility, Genetoul, anexplo, US006/INSERM, Toulouse, the Toulouse INSERM Metatoul-Lipidomique Core Facility-MetaboHub ANR-11-INBS-010, where lipidomic analysis were performed, and the platform Aninfimip, an EquipEx (‘Equipement d’Excellence’) supported by the French government through the Investments for the Future program (ANR-11-EQPX-0003). We do thank Fanny Lafouresse for critical reading of the manuscript.

## Author Contributions

Conceptualization: E.E., E.L. R.J. and N.C., Validation: E.L., E.E., R.J. and N.C; Formal Analysis: E.E, Investigation: E.L., E.E., R.J., C.P., G.P. and X.M.O; Resources: N.G. and C.L; Writing – Original draft: E.E., E.L., R.J., N.C., S.V., and G.D.; Visualization: E.E, N.C., C.P. and E.L.; Supervision E.E and N.C; Funding Acquisition: S.V., E.E., N.C. and G.D.

## Declaration of Interests

The authors declare no competing interest

## Notes

### Competing Interest Statement

The authors have declared no competing interest.

