## Supplementary material for "T helper cell-licensed mast cells promote inflammatory Th17 cells": Figure S

### Supplemental information

### Supplementary figures

**Figure S1**

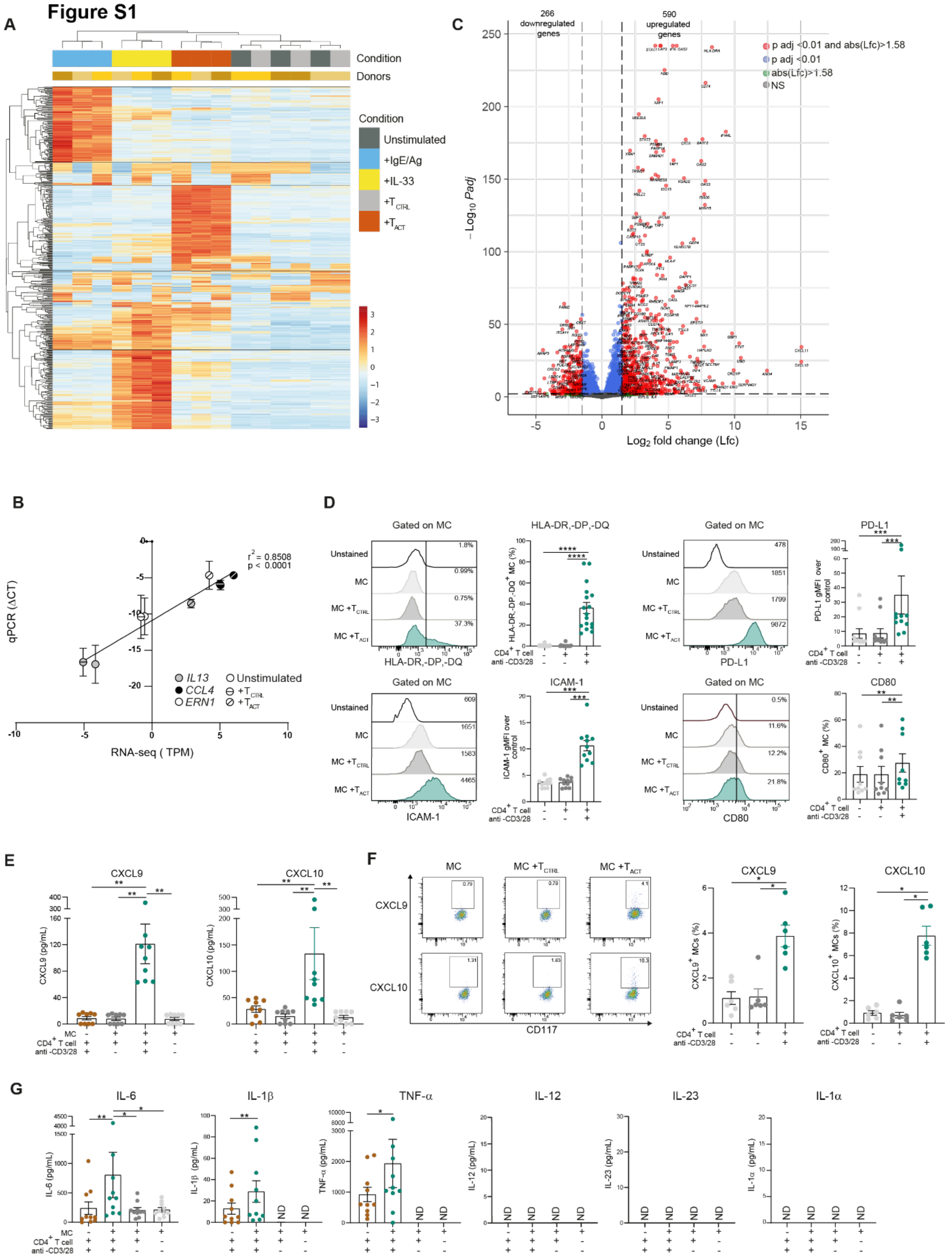

**Figure S1. mRNA and protein level analysis of MCs stimulated in indicated conditions. Related to Figure 1.**

(A) 500 most significantly differential expressed genes between the different conditions and control condition (Unstimulated MCs) were plotted in a heatmap. Regularized-logarithm transformed counts values (DESeq2, rlog function) were mean-centered and scaled for each gene. Hierarchical clustering of DEGs and samples was based on Euclidean distance using complete linkage function. Each column corresponds to one sample with indicated donor and stimulation condition and each row corresponds to a specific gene.

(B) Validation of RNA-seq data by RT-qPCR. *IL13, CCL4* and *ERN1* gene expression was measured by RT-qPCR from the same RNA extract used for RNA-seq. plotted are TPM versus $\Delta$CT values.

(C) Volcano plot showing the P-adjusted value (-log10-transformed) as a function of the fold-change (log2-transformed) from MC +$T_{ACT}$ relative to MC + $T_{CTRL}$. Genes with | log2(fold change) | > 1.58 and FDR < 0.01 (significantly differentially expressed genes) are in red, genes with | log2(fold change) | > 1.58 but FDR ≥ 0.01 are in green, genes with | log2(fold change) | ≤ 1.58 but FDR < 0.01 are in blue, and the rest are in gray.

(D) MCs were cocultured or not with $CD4^+$ T cells for 48 hours and HLA-DR,-DP,-DQ, ICAM-1, PD-L1, CD80 expression on MC surface was analyzed by flow cytometry. Representative histograms and pooled data (depicted as % positive cells for HLA-DR, DP, DQ and CD80 or geometric mean fluorescence intensity (gMFI) fold increase over the control for ICAM-1 and PD-L1), bars represent mean ± SEM; each point represents an experiment (n= 9-17).

(E-F) MCs were cocultured or not with $CD4^+$ T cells for 48h. CXCL9 or CXCL10 concentrations in the culture supernatant were measured by bead-based multiplex assay (n=9) (E). Representative dot plots of CXCL9 or CXCL10 intracellular staining gated on MCs ($CD117^+$ cells) and pooled data from 6 independent experiments; bars represent mean ± SEM; each point represents an experiment.

(G) MCs were cocultured or not with $CD4^+$ T cells for 48h. IL-6, IL1$\beta$, TNF, IL-12, IL-23 and IL-1$\alpha$ concentrations in the culture supernatant were measured by bead-based multiplex assay. Bars represent mean ± SEM; each point represents an experiment (n=10).

Friedman test followed by paired Wilcoxon signed-rank test. *p < 0.05, **p < 0.01, ***p < 0.001, ****p < 0.0001, ns not significant.

**Figure S2**

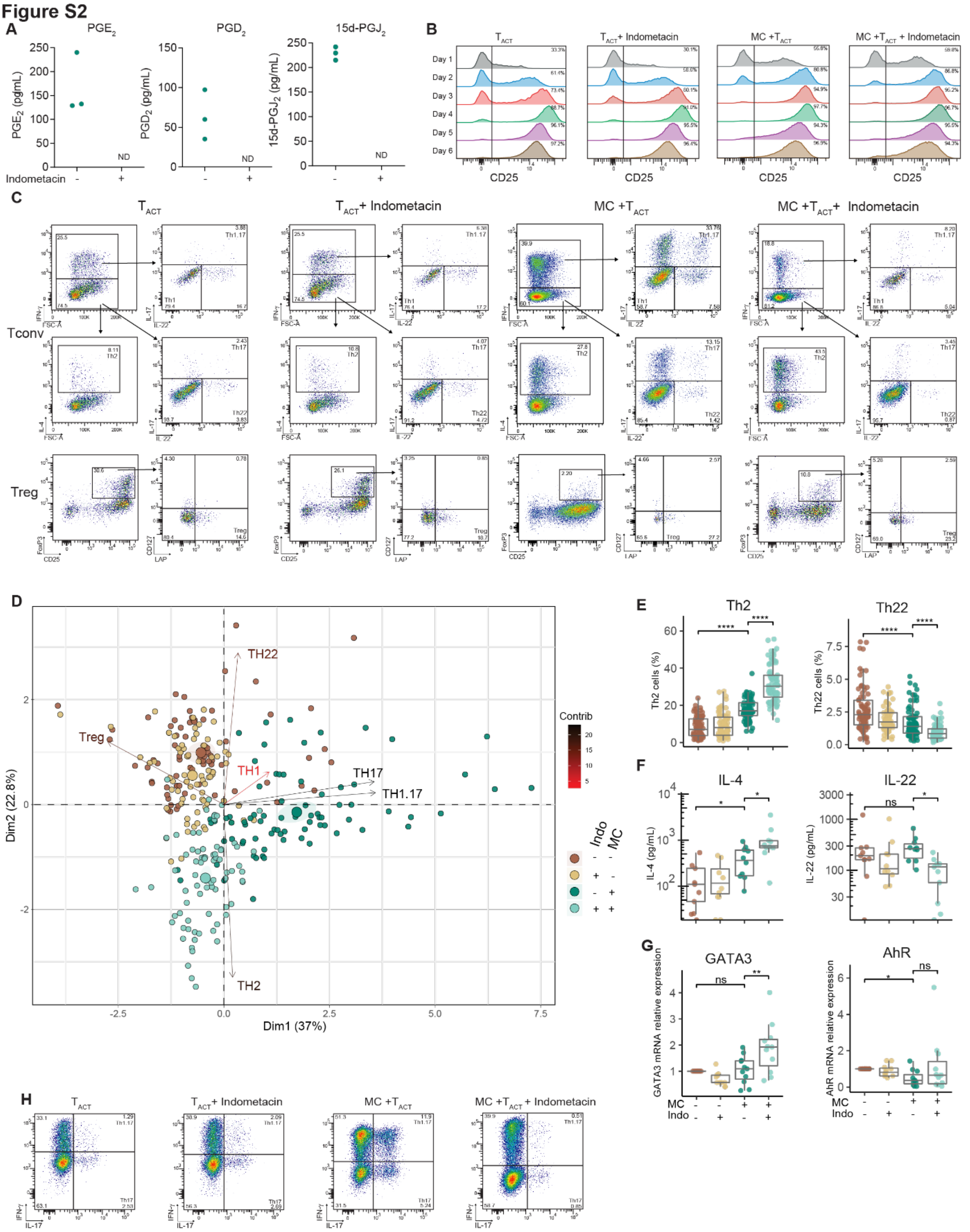


**Figure S2. MC$^{TH}$ drive Th cells toward IL-17 production in a COX-2 dependent manner. Related to Figure 3.**

(A) MCs were cocultured with CD4$^{+}$ T cells plus anti-CD3/28-coated beads for 48h with or without indometacin (100 µmol/L). Indicated eicosanoid concentration were measured by LC-MS/MS. Each point represents one MC/T cell pair from 3 different donors; ND, not detected.

Effector/memory CD4$^+$ T cells were cocultured with MCs in presence of anti-CD3/CD28-coated beads for 6 days with or without indometacin.

(B) Kinetics of CD25 expression in CD4$^+$ T cells (representative histograms, numbers indicate % CD25$^+$ cells).

(C) Th cell subsets (Tconv) and Treg analysis and gating strategy at day 6 of coculture (Representative dot plots of indicated conditions)

(D) Principal Component Analysis was performed using frequencies of identified subsets in each MC/T cell cocultures. Shown is a biplot that overlays individuals (each point represents a MC/T cell coculture, larger point represents the group centroids) and loadings (loading contribution is represented by color gradient)

(E) Frequencies of Th2 and Th22 cells are presented as box and whiskers plot (Tukey style), each point represents a MC/T cell pair. Pooled data (n=77) from 31 independent experiments, Friedman test and pairwise comparisons using Dunn's test.

(F-G) After 6 days coculture, CD4$^+$ T cells were FASC-sorted. IL-4 and IL-22 amount were measured after restimulation with PMA and ionomycin (n=9 from 3 independent experiments) (F). GATA3 and AhR expression was assessed by RT-qPCR (n=12 from 8 independent experiments) (G). Data are presented as box and whiskers plot (Tukey style), each point represents a MC/T cell pair. Friedman test and pairwise comparisons using paired Wilcoxon signed-rank test.

(H) Flow cytometry analysis of Th17 and Th1.17 subset frequencies at day 12 of coculture. Representative dot plots.

**Figure S3**

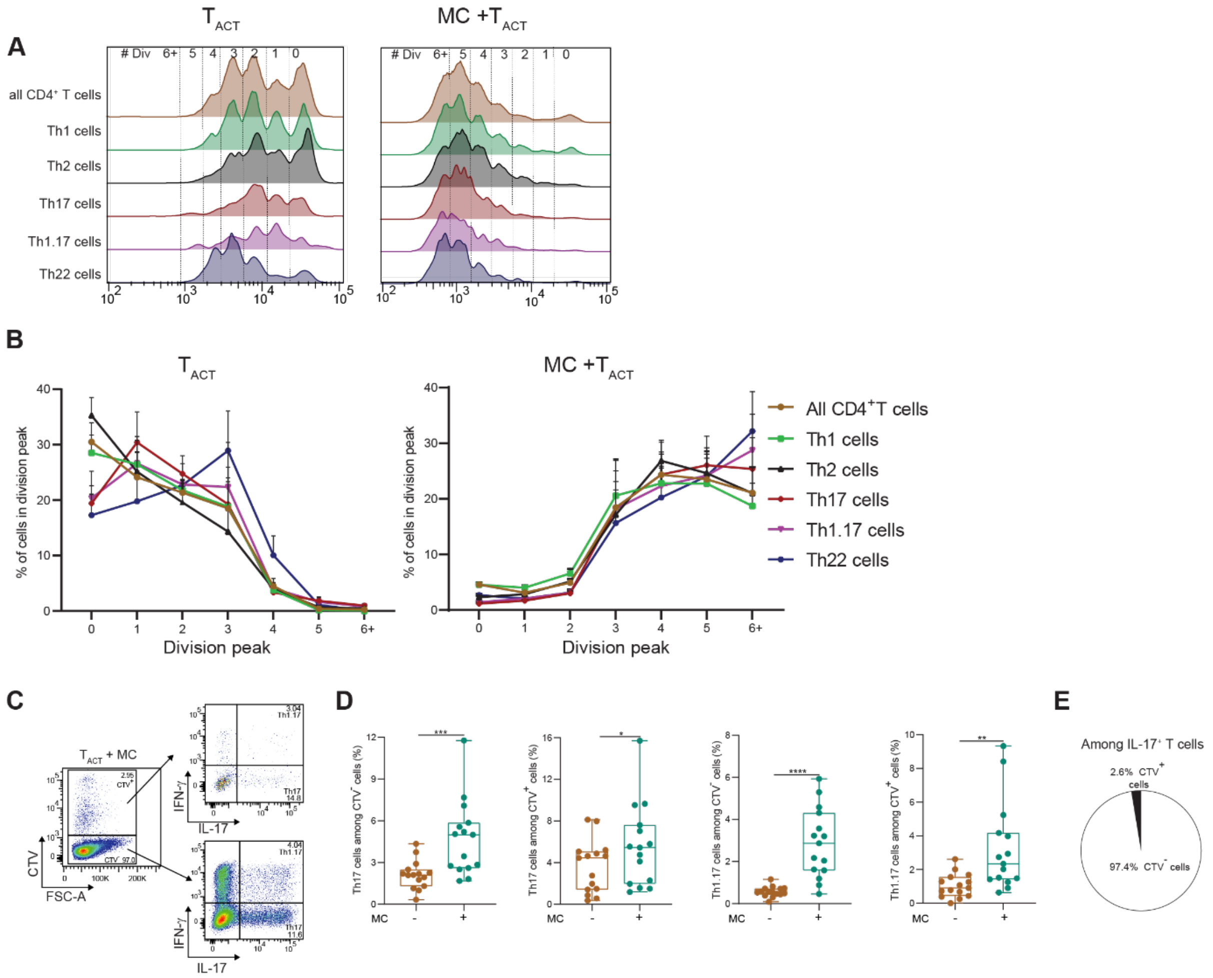


**Figure S3. Neither Th17 proliferative advantage nor Treg differentiation into Th17 cells account for Th17 emergence in presence of MC$^{TH}$. Related to Figure 4**

(A-B) CTV-labelled CD4$^{+}$ T cells were cocultured with MCs for 4 days and restimulated with PMA/ionomycin to detect intracellular cytokine produced. Representative histogram showing CTV dilution across the different subsets identified (A) and pooled data from 8 MC/T pairs from 4 independent experiments, proportion of each subset in each CTV division peak (mean ± SEM). One-way ANOVA followed by Tukey's multiple comparison test showed no significant differences among Th cell subsets.

(C-E) Sorted Treg cells were labelled with CTV cell tracer, reincorporated with their Tconv counterparts and next cocultured with MCs for 6 days. Representative IFN-γ and IL-17 dot plots (C) and pooled data from 10 independent experiments (D). Data are presented as box and whiskers plot; each point represents a MC/T cell pair (n=15). Proportion of Treg (CTV$^{+}$) and Tconv (CTV$^{-}$) cells among IL-17$^{+}$ T cells (E).

#### Figure S4

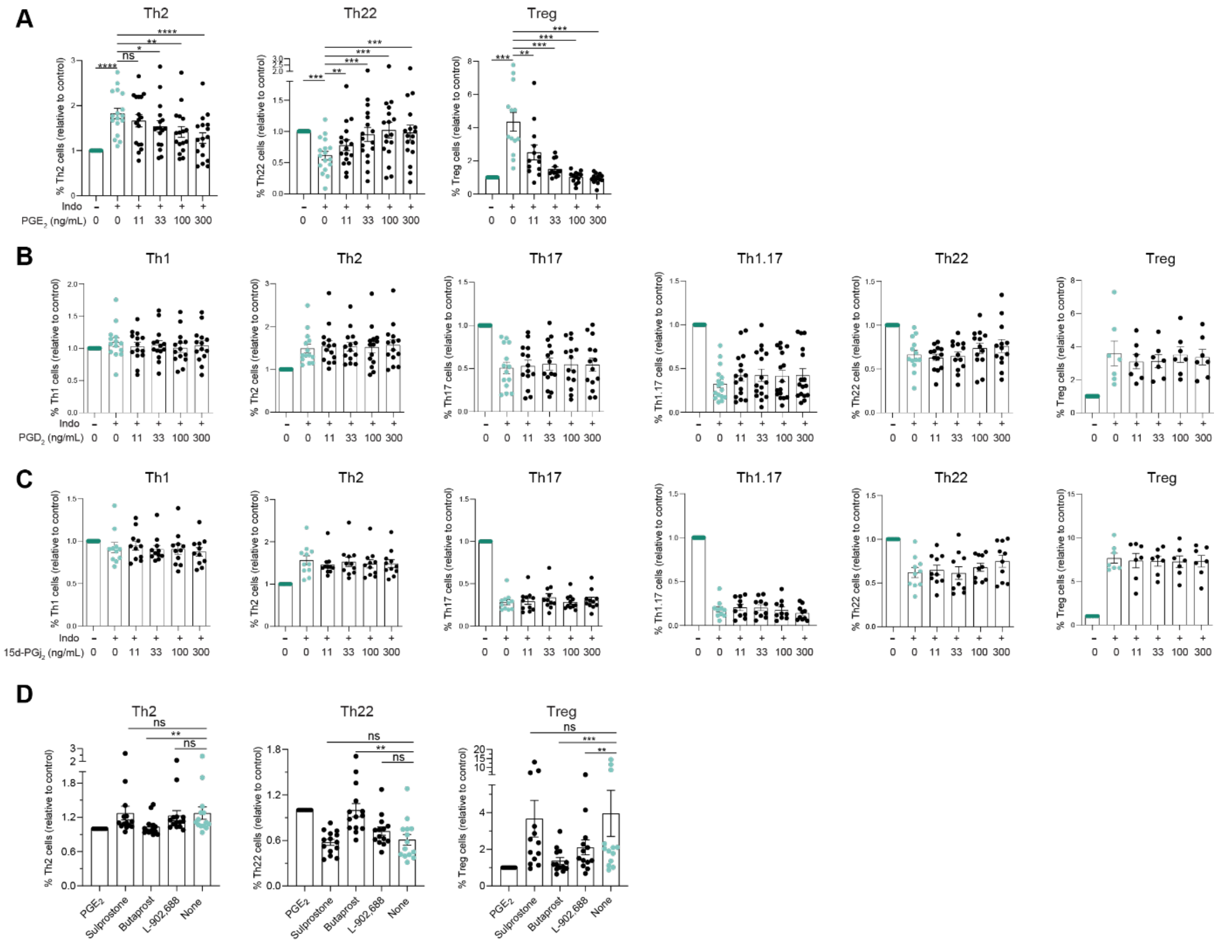


**Figure S4. Prostaglandin impact on Th cell subsets distribution. Related to Figure 5**

(A) Indometacin-treated cocultures were supplemented with increasing concentrations of $PGE_2$ and $CD4^+$ T cell subsets frequencies were analyzed by flow cytometry at day 6. $CD4^+$ T cell subset frequencies are presented as fold change over the control condition (MC+ $T_{ACT}$), mean ± SEM, each point represents an experiment (n=17 from 13 independent experiments)

(B-C) Effector/memory $CD4^+$ T cells were cocultured with MCs in presence of anti-CD3/CD28 coated beads for 6 days. Indometacin-treated cocultures were supplemented with increasing concentrations of $PGD_2$ (B) or 15d-$PGJ_2$ (C) and $CD4^+$ T cell subsets frequencies were analyzed by flow cytometry at day 6. Bars represent mean ± SEM (n=10-16). ANOVA showed no significant difference among groups.

(D) Indometacin-treated cocultures were supplemented with $PGE_2$ receptor agonists or 300 ng/mL of $PGE_2$. Th cell subsets frequencies were analyzed by flow cytometry at day 6. $CD4^+$ T cell subset frequencies are presented as fold change over the control condition ($T_{ACT}$ + MC with indometacin and 300 ng/mL $PGE_2$), mean ± SEM, each point represents an experiment (n=14 from 4 independent experiments).

#### Supplementary Table legends

| Transcription factors | NES | # Targets | # Motifs/Tracks |
|---|---|---|---|
| **+ TACT** | | | |
| IRF9, IRF8, IRF4, IRF7, IRF1, IRF2, IRF5, IRF3, IRF6 | 16.46 | 606 | 8 |
| STAT1, STAT2 | 14.29 | 704 | 29 |
| STAT2 | 9.62 | 130 | 2 |
| IRF1 | 8.21 | 154 | 1 |
| STAT1 | 7.91 | 231 | 5 |
| IRF2 | 6.75 | 459 | 4 |
| IRF4 | 6.38 | 376 | 4 |
| BCL3 | 6.15 | 564 | 30 |
| IRF8 | 4.29 | 131 | 3 |
| ATF3 | 3.68 | 288 | 3 |
| NFKB1 | 3.61 | 357 | 5 |
| STAT3 | 3.29 | 79 | 1 |
| **+ IgE/Ag** | | | |
| BATF, FOS, FOSB, FOSL1, JUNB, JUND, JUN, BACH2 | 5.43 | 923 | 51 |
| MYC | 4.61 | 475 | 5 |
| SRF | 4.25 | 195 | 6 |
| ATF3, CREB1, CREM, ATF2, ATF1, ATF4, ATF6, ATF7 | 4.05 | 956 | 30 |
| MAX | 3.59 | 460 | 4 |
| NFKB1, NFKB2, BCL3, RELB, RELA, OVOL2, EBF1, STAT6 | 3.48 | 184 | 5 |
| RARB | 3.45 | 447 | 1 |
| MXI1 | 3.45 | 286 | 1 |
| FOSL2 | 3.29 | 207 | 2 |
| XBP1, CREB3 | 3.17 | 91 | 2 |
| **+ IL-33** | | | |
| BCL3, NFKB1, NFKB2, RELB, RELA, REL | 7.96 | 811 | 81 |
| NFKB1 | 5.65 | 448 | 9 |

| | | | |
|---|---|---|---|
| IRF4, SPI1, E2F1, MYB | 3.55 | 157 | 2 |
| MZF1, PURA | 3.12 | 66 | 1 |
| NFIC | 3.01 | 199 | 1 |

**Table S1. Master regulator analysis via iRegulon. Related to Figure 1**

MC upregulated genes (FDR<0.01 and fold change>2) in indicated stimulation conditions were analyzed by iRegulon to identify enriched transcription factor binding, based on the TRANSFAC database and ENCODE (using iRegulon). Only transcription factor motifs with normalized enrichment scores (NES) > 3 (indicating a motif that covers a large proportion of the input genes) are listed. Upregulated transcription factors in indicated stimulation condition as compared to unstimulated MCs (FDR<0.01 and fold change>2) are underlined.

**Table S2. DEG list of $MC^{TH}$**

DEGs from $MC+T_{ACT}$ relative to $MC+T_{CTRL}$ condition (FDR<0.01 and $|Log_2foldfchange|>1.58$). Related to Figure 1.

**Table S3 Ligand-Receptor interaction inference**

predicted ligands from $MC^{TH}$ (only ligands corresponding to DEGs from $MC+T_{ACT}$ relative to $MC+T_{CTRL}$ were analyzed) with their associated receptors expressed on activated memory $CD4^+$ T cells. the last column provides the communication score. Related to figure 1.

| Gene (Ensembl) | Gene (symbol) |
|---|---|
| ENSG00000239713 | **APOBEC3G** |
| ENSG00000115009 | **CCL20** |
| ENSG00000121807 | **CCR2** |
| ENSG00000101017 | **CD40** |
| ENSG00000013725 | **CD6** |
| ENSG00000164400 | **CSF2** |
| ENSG00000163599 | **CTLA4** |
| ENSG00000169429 | **CXCL8** |
| ENSG00000088305 | **DNMT3B** |
| ENSG00000164308 | **ERAP2** |
| ENSG00000204525 | **HLA-C** |
| ENSG00000196735 | **HLA-DQA1** |
| ENSG00000179344 | **HLA-DQB1** |
| ENSG00000204287 | **HLA-DRA** |

| | |
|---|---|
| ENSG00000196126 | **HLA-DRB1** |
| ENSG00000090339 | **ICAM1** |
| ENSG00000163600 | **ICOS** |
| ENSG00000115267 | **IFIH1** |
| ENSG00000111537 | **IFNG** |
| ENSG00000136634 | **IL10** |
| ENSG00000169194 | **IL13** |
| ENSG00000134470 | **IL15RA** |
| ENSG00000115607 | **IL18RAP** |
| ENSG00000134460 | **IL2RA** |
| ENSG00000164399 | **IL3** |
| ENSG00000113525 | **IL5** |
| ENSG00000125347 | **IRF1** |
| ENSG00000137265 | **IRF4** |
| ENSG00000140968 | **IRF8** |
| ENSG00000168961 | **LGALS9** |
| ENSG00000113594 | **LIFR** |
| ENSG00000107968 | **MAP3K8** |
| ENSG00000165030 | **NFIL3** |
| ENSG00000109320 | **NFKB1** |
| ENSG00000077150 | **NFKB2** |
| ENSG00000144802 | **NFKBIZ** |
| ENSG00000167207 | **NOD2** |
| ENSG00000099985 | **OSM** |
| ENSG00000117090 | **SLAMF1** |
| ENSG00000185338 | **SOCS1** |
| ENSG00000115415 | **STAT1** |
| ENSG00000138378 | **STAT4** |
| ENSG00000164691 | **TAGAP** |
| ENSG00000049249 | **TNFRSF9** |
| ENSG00000106952 | **TNFSF8** |

**Table S4. Overlapping genes between prioritized gene candidates identified in IBD GWAS and upregulated genes in MC$^{TH}$.** Related to Figure 6.
